# Functional genomics reveals an off-target dependency of drug synergy in gastric cancer therapy

**DOI:** 10.1101/2023.10.07.561351

**Authors:** Ozen Leylek, Megan E. Honeywell, Michael J. Lee, Michael T. Hemann, Gulnihal Ozcan

## Abstract

The rational combination of anticancer agents is critical to improving patient outcomes in cancer. Nonetheless, most combination regimens in the clinic result from empirical methodologies disregarding insight into the mechanism of action and missing the opportunity to improve therapy outcomes incrementally. Deciphering the genetic dependencies and vulnerabilities responsible for synergistic interactions is crucial for rationally developing effective anticancer drug combinations. Hence, we screened pairwise pharmacological interactions between molecular-targeted agents and conventional chemotherapeutics and examined the genome-scale genetic dependencies in gastric adenocarcinoma cell models. Since this type of cancer is mainly chemoresistant and incurable, clinical situations demand effective combination strategies. Our pairwise combination screen revealed SN38/erlotinib as the drug pair with the most robust synergism. Genome-wide CRISPR screening and a shRNA-based signature assay indicated that the genetic dependency/vulnerability signature of SN38/erlotinib is the same as SN38 alone. Additional investigation revealed that the enhanced cell death with improved death kinetics caused by the SN38/erlotinib combination is surprisingly due to erlotinib’s off-target effect that inhibits ABCG2 but not its on-target effect on EGFR. Our results confirm that a genetic dependency signature different from the single-drug application may not be necessary for the synergistic interaction of molecular-targeted agents with conventional chemotherapeutics in gastric adenocarcinoma. The findings also demonstrated the efficacy of functional genomics approaches in unveiling biologically validated mechanisms of pharmacological interactions.

**Significance:** Functional genomics approaches efficiently demonstrated an off-target dependency of the synergistic interaction of erlotinib with SN38 in gastric adenocarcinoma cell models.

## Introduction

Targeting key oncogenic drivers and signaling mechanisms by molecular-targeted agents is becoming an attractive strategy to increase sensitivity to chemotherapy in cancer, replacing earlier regimens of multiple chemotherapeutics. The non-overlapping modes of actions of molecular-targeted agents with conventional chemotherapeutics can address tumor heterogeneity, prevent cross-resistance, and achieve a more tumor-specific activity (1,2). However, distinct modes of action do not guarantee a synergistic effect or, at least, an additive effect on cancer cell death (3). A rational selection of the combinatorial agents based on a clear understanding of the genetic dependencies and vulnerabilities for synergistic action is valuable to improve patient outcomes while minimizing the risk of treatment failure from empirical combination approaches (4,5). Such an understanding also paves the way for discovering novel agents with potent synergy in anticancer action.

Functional genomics screens have emerged as powerful tools for unveiling the molecular mechanisms of drug action and pharmacological interactions in combination therapy. Such screens have revealed critical dependencies for drug response and chemoresistance successively in cancer cells. For instance, CRISPR screening and clone tracing showed low cross-resistance among the constituents of the R-CHOP (rituximab, cyclophosphamide, adriamycin, oncovin, prednisolone) regimen (6). Pritchard and colleagues developed a shRNA-based signature assay to characterize and predict the mechanisms of drug action, which enabled the dissection of the mode of synergism (5,7). Integrating shRNA-based signature assays with informatics tools, they showed that synergistic chemotherapeutics can exhibit a genetic signature identical to the signature of the dominant chemotherapeutic within the combination, as in the case of 5-FU/Leucovorin, where leucovorin reinforces the action of 5-FU. The authors also showed that some other combinations may exhibit an average of the genetic dependencies of individual chemotherapeutics in the combination, exemplified by actinomycin D/chlorambucil and CHOP. When combined, drug molecules may also reveal a novel genetic signature distinct from their signatures as monotherapy, as suggested in yeast models (8). In theory, molecular-targeted agents and conventional chemotherapeutics may also interact in line with one of these genetic dependency profiles when combined. However, in the context of targeted agents, understanding the mechanisms underlying drug synergy is particularly challenging, as these drugs can reprogram intracellular signaling pathways, a mode of action distinct from conventional chemotherapeutics. Moreover, emerging evidence indicates that the anti-cancer efficacy of many molecular-targeted agents is dependent on off-targets rather than known primary targets (9). Hence, we investigated the genetic dependency signature of synergistic molecular-targeted agent/conventional chemotherapeutic pairs using functional genomics approaches in gastric adenocarcinoma.

Gastric adenocarcinoma is an intractable and chemo-resistant cancer with only a few molecular-targeted agents available for a limited number of patients (10). Several molecular targets, such as epidermal growth factor receptor (EGFR), mammalian target of rapamycin (mTOR), and c-met, have significant involvement in the aggressiveness of gastric adenocarcinomas. However, drugs targeting these proteins have not achieved the clinical success promised in preclinical studies and have yet to be translated into the clinic (11–13). This failure may result from an irrational combination of these agents with other anticancer agents. Therefore, combining EGFR, mTOR, and c-met inhibitors with chemotherapeutics based on a logic of pharmacological and genetic interactions is pivotal to therapeutic success.

In this study, we screened pairwise pharmacological interactions of EGFR, mTOR, and c-met inhibitors with chemotherapeutics from five distinct groups in gastric adenocarcinoma cells, rigorously inspecting the kinetics of cell death. To understand the mechanisms of synergy, we investigated the genetic dependency signatures using a genome-wide CRISPR screen and the shRNA-based signature assay.

## Materials and Methods

### Cell lines and culture conditions

AGS (American Tissue Type Culture Collection, USA), SNU1, SNU5, SNU16, SNU484, and NCI-N87 gastric adenocarcinoma cell lines (Korean Cell Line Bank), and Lenti-X cells were grown in RPMI (Corning, USA) supplemented with 10% fetal bovine serum (FBS) (Gibco, USA) and 1% Penicillin-Streptomycin (P/S) (Gibco, USA). Eμ-Myc Cdkn2a^Arf−/−^ leukemia cell line was grown in B-cell media (BCM) composed of DMEM and IMDM media supplemented with 10% FBS, 1% P/S and 0.1% 2-mercaptoethanol (Gibco, USA).

### Assessment of pharmacological interactions for drug pairs

We analyzed drug responses with MTT assay at the initial screening. Briefly, we treated the cells in 96-well cell culture plates (Corning, USA) with at least seven different concentrations of each drug or the drug combinations. After five days of incubation, 3 mg/ml MTT was added to each well and incubated for 4 hours in the dark. Formazan crystals, the reduced form of MTT, were dissolved with 13.8% SDS solution (Bio-Rad, USA) containing 1% HCl on a shaker overnight at room temperature, and the absorbance was measured at 570 nm by a microplate reader (Synergy H1, Biotek). The relative viability (RV), the ratio of the signal from live cells in each treated group to the control group, was calculated. We plotted each treatment’s sigmoidal dose-response curve (DRC) using four-parameter logistic regression to assess plateau, hill slope, EC_max_, and EC50 in GraphPad Prism7. In the next step, we inferred the fraction-affected (fa) values normalized to 1 from RV. We computed the combination indices (CI) using fa values for each concentration of monotherapy and combinations on the Compusyn program developed by Chou and Talalay (14). CI<1, CI=1, and CI>1 denote synergism, additivity, and antagonism, respectively, for drug pairs.

### Fluorescence-based and lysis-dependent inference of cell death kinetics assay

We performed the fluorescence-based and lysis-dependent inference of cell death kinetics (FLICK) assay to investigate drug-induced cell death, as described in Richard et al. (15). Shortly, we seeded the cells at a density of 2000-2500 cells/well in 96-well plates. The following day, we treated cells with the drug of interest. We used the SYTOX dye (Invitrogen, USA) to indicate dead cells. The fluorescence of SYTOX was monitored throughout the assay using Tecan Spark or Tecan M1000 plate readers to infer the cell death kinetics. At the end of the experiment, we exposed the cells to 0.1% Triton-X (Thermo Scientific, USA) for lysis, from which we deduced the total population size through the SYTOX signal. In addition to experiment plates, we prepared an untreated ’*T0*’ plate for the lysis of the cells at the beginning of the experiment to determine the initial cell population size. We used the exponential growth modeling of population sizes at the initial and end time points to extrapolate the total population size at intermediate time points. We calculated the size of live populations using dead population sizes represented by the SYTOX signal and whole population sizes. Based on the information on live, dead, and total populations, we evaluated three different pharmacological metrics: fractional viability (FV), lethal fraction (LF), and normalized growth rate inhibition (GR). We provide the equations for each metric below. The code we used to perform curve fitting on MATLAB is available on GitHub (https://github.com/MJLee-Lab) (16).

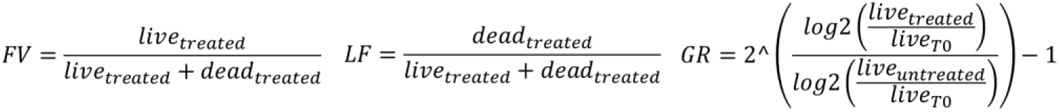

### Quantitative Immunoblotting

We washed the cell pellets with ice-cold PBS and lysed them in SDS-lysis buffer (50 mM Tris-HCl, 2% SDS, 5% glycerol, 5mM EDTA, 1mM NaF, 10 mM 𝛽-GP, 1mM phenylmethylsulphonyl-fluoride, 1mM Na_3_VO_4_, protease and phosphatase inhibitors). We loaded the lysates at equal protein amounts and concentrations on 4-15% precast TGX gels (Bio-Rad, USA) or 10% hand-poured SDS-PAGE gels. Proteins were transferred to nitrocellulose membranes, then blocked in 1:1 PBS/Intercept Blocking Buffer (IBB, LI-COR Biosciences, USA). Membranes were incubated with 1:1000 diluted primary antibodies at 4°C overnight. The next day, after washing, membranes were incubated with 1:15000 diluted secondary antibodies (LI-COR, USA) at room temperature for 1 hour. We visualized the immunoblots on the LI-COR Odyssey CLx scanner. Band intensities were quantified using ImageJ and normalized by internal loading control.

### Flow cytometry

Cell pellets were washed in ice-cold PBS, fixed in ice-cold 70% ethanol, and stored at -20°C. Before cell cycle progression analysis, each sample was washed twice in ice-cold PBS and resuspended in 10% RNase A in PBS. Propidium iodide (final concentration: 0.5mg/ml) was added to each sample. To assess apoptotic activity, cells were fixed in 4% formaldehyde in PBS at room temperature for 15 mins. After washing with cold PBS, samples were fixed in ice-cold 100% methanol and stored at -20°C. Before analysis, methanol was removed, and pellets were washed twice with PBS containing 0.1% Tween-20 (PBS-T). Each sample was incubated with 1:500 diluted primary cleaved-caspase3 antibody in 1:1 PBS/IBB at room temperature. After 8 hours, samples were washed with PBS-T and incubated with primary cleaved-PARP Alexa-647 antibody and goat anti-rabbit Alexa-488 (diluted 1:250 in 1:1 PBS-T/IBB) overnight at room temperature.

For DNA damage analysis, cells were incubated with primary phospho-H2AX antibody (diluted 1:200 in 1:1 PBS-T/IBB) at room temperature for 8 hours. Samples were washed with PBS-T and incubated with 1:250 diluted goat anti-mouse Alexa-488 overnight at room temperature. Then, samples were washed with PBS and resuspended in 10% RNase A in PBS. Propidium iodide was added to each sample (the final concentration was 0.5mg/ml). The samples were analyzed in LSRII or Miltenyi MACS Quant VYB flow cytometers. Data analysis was performed on FlowJo.

### Generation of SNU5-Cas9 and SNU5^EGFR-KO^ cells

pCMV-VSV-G (Addgene, USA) (17) and psPAX2 (a gift from Didier Trono; Addgene plasmid # 12260; Addgene, USA) packaging vectors and the equimolar amount of lentiCas9-Blast (Addgene, USA) (18) were diluted in OptiMEM. After adding lipofectamine 2000 and incubating at room temperature for 15 minutes, the mixture was added to lenti-X cells. After 6 hours, the medium was replaced with a fresh medium. The media with lentiviral particles were collected at 24 and 48 hours, then centrifuged and filtered with a 0.45μm filter. SNU5 parental cells were transduced with viruses containing lentiCas9-Blast by centrifuging at 830xg at 32°C for 90 minutes in the presence of polybrene (final concentration: 4ug/ml), which was followed by the replacement with fresh media. Cells expressing Cas9 (SNU5-Cas9) were selected by blasticidin (10μg/ml). SgRNA targeting EGFR (5’-CGATCTCCACATCCTGCCGG-3’) was cloned into the lentiGuide-puro vector (Addgene, USA) (18). SNU5-Cas9 cells were then infected with viruses containing this vector following the protocol described above. SNU5-Cas9 cells expressing sgRNA were selected by puromycin (2μg/ml). Single-cell clones were generated by seeding cells into 96-well plates, with one cell per well. The EGFR expression in each clone was evaluated via immunoblotting. The gene knockout was further confirmed for the selected SNU5EGFR-KO clone using Sanger sequencing.

### Genome-wide CRISPR Screen

We performed a genome-wide CRISPR screening using Toronto KnockOut Library v3 (TKOv3) with 71090 sgRNAs (70948 sgRNA targeting 18053 protein-coding genes - 4 sgRNA/gene - and 142 control non-targeting sgRNAs against EGFP, LacZ, and luciferase) (Addgene, USA) (19). More than 300x10^6^ SNU5-Cas9 cells were infected with the library to achieve 500x coverage. After expansion in fresh media, the cells were plated in duplicate 90x10^6^ and 50x10^6^ cells for each treatment and untreated condition, respectively. 50x10^6^ cells were collected and frozen in duplicate for the T0 controls. Cells were treated with either SN38 (13.5nM) or the combination (3.7nM SN38 + 370nM Erlotinib) on day 0. On day 2, cells in all experiment groups were passaged by maintaining culture conditions. Live and dead fractions in each group were separated using annexin V-conjugated magnetic bead sorting on day 3.

Genomic DNA from samples was extracted with the genomic DNA purification kit (Promega, USA). The TKOv3 CRISPR sequencing library was prepared using a two-step PCR reaction: the PCR1 to enrich the gRNA regions in the genome and the PCR2 to amplify gRNAs with sequencing adapters. The 200bp PCR2 product was excised, and DNA was purified from agarose gel using a gel extraction kit (Qiagen, Germany). Reads of each sequenced library were demultiplexed using the barcode splitter and trimmer functions of the FASTX toolkit. Reads were mapped to the TKOv3 library using the Bowtie2 read alignment tool. A parametric fit of DESeq2 was used to determine the sgRNA-level log2-fold change (L2FC) for each comparison of interest. Then, we randomly assigned the nontargeting sgRNA controls to 36 sets of four sgRNAs per gene. Guide-level scores were transformed into the single gene-level fold change (FC) by calculating the mean of all sgRNAs. FCs at the gene level were *z*-scored based on the mean and standard deviation of nontargeting genes. Empiric p-values were calculated from *z*-scored FCs by bootstrapping guide-level scores, which were then false discovery rate (FDR)-corrected using the Benjamini-Hochberg procedure.

### CRISPR Screen Validation

All sgRNAs targeting selected candidate hits and nontargeting control sgRNA targeting the LacZ gene were cloned in the pRDA_170 vector with the e2-crimson dark red fluorescence protein sequence. SNU5-Cas9 cells were transduced with viruses containing corresponding vectors to generate the knockout bulk populations (SNU5*^Gene^ ^X^*^-KO^) and the untargeted control population (SNU5^LacZ^), followed by puromycin selection. Two different approaches were adopted for the validation, and the treatments were performed at doses applied in the genomic screening. The first was done by FLICK assay, as explained above. The second validation was performed using a competition assay. For this, SNU5-Cas9 and SNU5-Cas9 cells with sgRNA targeting gene of interest were mixed at a 1:1 ratio and seeded in 12-well plates (2x10^5^ cells/well). The enrichment/depletion of e2-crimson-positive cells was assessed after three days of drug treatment using BD FACS Celesta flow cytometry. Data analysis was performed on FlowJo.

### RNAi-based Signature Assay

Eight shRNAs in pMSCV-LTR-miR30-SV40-GFP (MLS) retroviral vector, validated for their knock-down and off-target effects, were used for signature assay (5,7). Eμ-Myc Cdkn2a^Arf−/−^ cells were transduced with each GFP-tagged shRNA construct at a 25-30% infection rate. Cells were seeded into 24-well plates in 250μl BCM and treated with 250 μl of drug-containing media, inducing 80-90% cell death (LD80-90) at 48 hours. At the endpoint, the live cells were examined by DAPI exclusion, and the percentage of GFP-positive cells was determined using BD FACS Celesta flow cytometry. Resistance index (RI), calculated following the formula: (GFPtreatment – (GFPtreatment* GFPcontrol))/ (GFPcontrol – (GFPtreatment* GFPcontrol)), was used to generate the signature of each drug of interest. Each signature was then compared with a previously established reference set of drugs using the modified Euclidean K-nearest neighbors algorithm via MATLAB, generating linkage ratios (LR) and p-values (20). An LR≤1 and a p-value <0.05 indicates a significant classification of the drug of interest into a specific category in the reference set. LR and/or p-values outside of these cut-offs are interpreted as the drug of interest that belongs to a "new class" with a mechanism of action not represented in the reference set.

### qRT-PCR

RNA was extracted from 1-1.5x10^6^ cells using the Bio-Rad RNA extraction kit (USA). The RNA was combined with 2.5μl of 2nM dNTPs (Life Technologies, USA) and 1μl of random hexamers (Invitrogen, USA) per sample for cDNA synthesis. After incubation at 65°C for 5 mins, an RT mix of 5μl first-strand buffer (Invitrogen, USA), 2μl of 0.1M DTT (Invitrogen, USA), and 0.5μl of Rnasin (Promega, USA) were added to each tube and incubated at room temperature for 10mins. Finally, 1μl of MMLV-RT enzyme (Invitrogen, USA) was added. The reactions were incubated at 37°C for 1h and then inactivated at 70°C for 15 mins. For the qRT-PCR reaction, 10μl of 2X Fast SYBR Green Master Mix (Applied Biosystems, USA), 1μl of 4μM primer mix (ABCG2-F: AGCCACAGAGATCATAGAGCC, ABCG2-R: TTCTCACCCCCGGAAAGTTG, GAPDH-F: AGCCACATCGCTCAGACAC, GAPDH-R: GCCCAATACGACCAAATCC) and 7μl nuclease-free grade water were mixed with 2μl of diluted cDNA sample in each well of a 96-well optical plate (Applied Biosystems, USA). The CT values for each reaction were determined using StepOnePlus Real-Time PCR System (Applied Biosystems). The GAPDH housekeeping gene was used to calculate the relative expression level of ABCG2.

### Assessment of Efflux Pump Activity

SNU5 cells seeded in 12-well plates were treated with erlotinib, osimertinib, and ABCG2i-1, ABCG2i-2 at 10, 1, 0.1, and 0.01μM. The following day, cells were exposed to Hoechst-33342 (final concentration: 5μg/mL) for 45 minutes at 37°C in the dark. Cells were washed with cold Hanks balanced saline solution (Gibco, USA) with 1mM HEPES (Gibco, USA) and 2% FBS (HBSS+), then taken into fresh media. Hoechst-33342 negativity was evaluated at times 0, 1h, and 16h using the FACSymphony A3 flow cytometer (BD Biosciences, USA) by analyzing at least 10,000 events per sample. Data analyses were performed on FlowJo.

## Results

### Identifying synergistic molecular-targeted agent and conventional chemotherapeutic pairs in gastric adenocarcinoma cells

To identify molecular-targeted agent-conventional chemotherapeutic pairs with strong synergism, we combined EGFR inhibitor erlotinib, mTOR inhibitor everolimus, and c-met inhibitor JNJ-38877605 with chemotherapeutics from five distinct groups acting via diverse mechanisms: doxorubicin (anthracycline), cisplatin (platinum derivative), 5-fluorouracil (fluoropyrimidine), paclitaxel (taxane), and SN38 (topoisomerase I inhibitor) (Figure 1A). We investigated the pharmacological interactions for these 15 drug pairs in AGS, SNU1, SNU5, and SNU16 gastric adenocarcinoma cell lines using Chou-Talalay’s CI method (Figure 1B, Figure S1A) (14).

**Figure 1.**
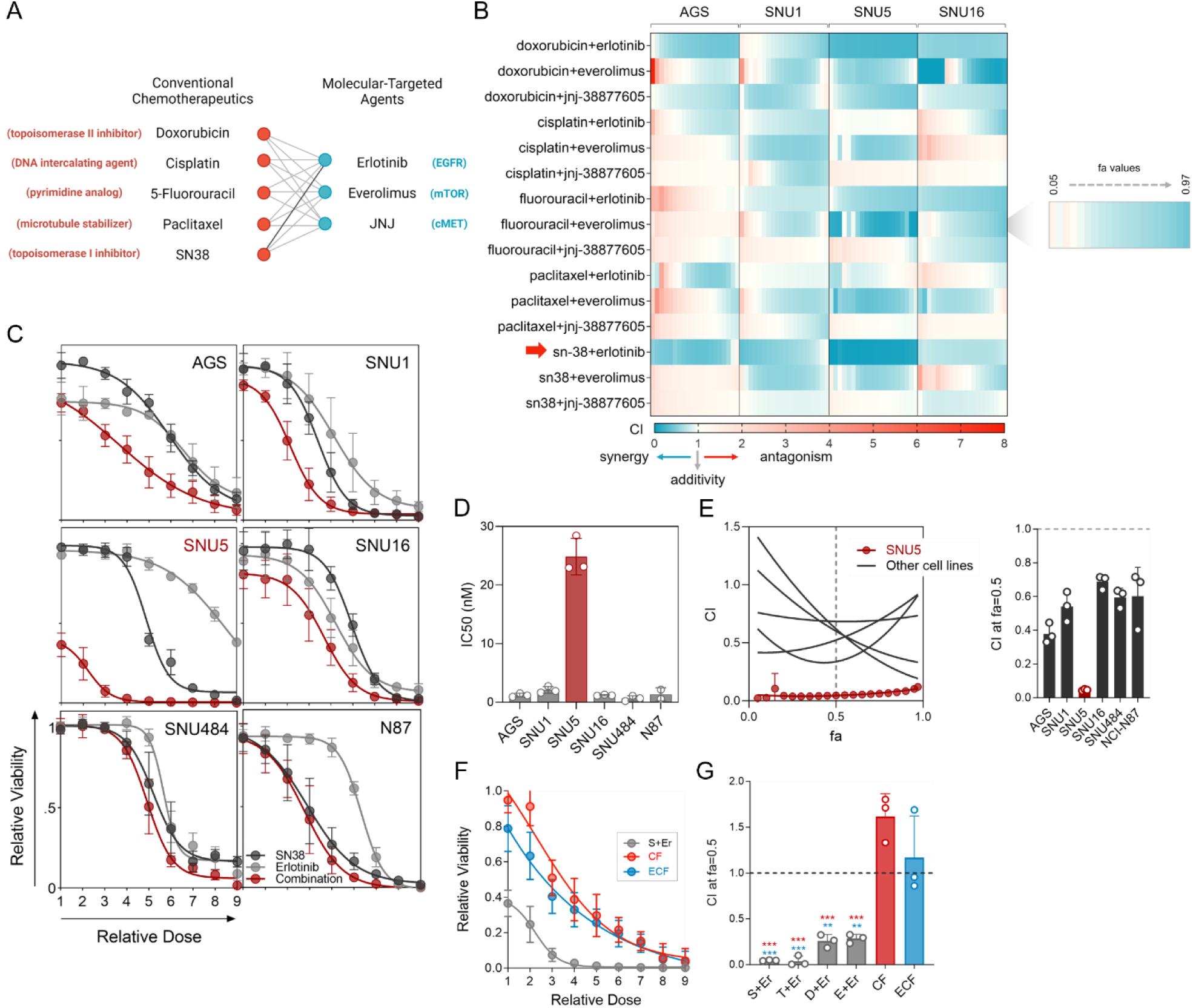
Pharmacological interactions for the molecular-targeted agent and conventional chemotherapeutic combinations in gastric adenocarcinoma cells. **(A)** The molecular-targeted agent and conventional chemotherapeutic pairs assessed in this study. The figure was generated on BioRender. (**B)** Heat map of combination indices (CI) at fraction affected (fa) values ranging from 0.05-0.97 for each drug pair in AGS, SNU1, SNU5, and SNU16 cells calculated by Chou-Talalay’s method. Blue indicates synergism, and white and red indicate additivity and antagonism. The strongest synergism was observed for the SN38-erlotinib pair in SNU5 cells (highlighted by a red arrow). (**C)** Dose-response curves for SN38, erlotinib, and SN38/erlotinib combination in AGS, SNU1, SNU5, SNU16, SNU484, and NCI-N87 gastric adenocarcinoma cells. **(D)** IC50 values for SN38 in all cell lines. **(E)** CI-fa plots generated using the data in C. **(F)** Dose-response curves for SN38/erlotinib, Cisplatin-5-Fluorouracil (CF), and Epirubicin-Cisplatin-Fluorouracil (ECF) combinations in SNU5 cells. **(G)** The CI’s for the combination of erlotinib with topoisomerase poisons (SN38, topotecan, epirubicin, and doxorubicin), CF, and ECF regimens (left). The CI plots were generated using the data in C, F, and Figures S1-C and D. Dunnett’s multiple comparisons test was used for statistical analysis. *** *p*<0.001, ** *p*=0.001.

The combination of erlotinib with SN38, the active metabolite of irinotecan, emerged as the most synergistic combination in the pairwise pharmacological interaction screen (Figure 1B). The potency of the SN38/erlotinib combination was significantly higher than that of SN38 and erlotinib as single agents in six different gastric adenocarcinoma cell lines (Figure 1C). Among these cell lines, SNU5 cells, derived from the metastatic ascites fluid of a poorly differentiated gastric adenocarcinoma patient who had previously undergone chemotherapy (21), exhibited the lowest sensitivity to SN38 (Figure 1D). Despite that, the synergism for SN38/erlotinib combination was the strongest in this cell line (Figure 1E, Figure S1B). The SN38/erlotinib combination was much more potent than the cisplatin/5-FU (CF) and epirubicin/cisplatin/5-FU (ECF) combinations, commonly used chemotherapy regimens in gastric cancer therapy (Figure 1F) (22). To interrogate whether the synergistic interaction of erlotinib with SN38 is unique to SN38 or shared by other topoisomerase inhibitors, we tested its combination with another topoisomerase I inhibitor, topotecan, and topoisomerase II inhibitors, doxorubicin and epirubicin in SNU5 cells (Figure 1G, Figure S1C-D). The combination of erlotinib with all topoisomerase poisons exhibited robust synergism in contrast to the antagonistic interaction in CF and ECF. However, the degree of synergism was highest for SN38/erlotinib combination.

### Validating the synergism in SN38/erlotinib combination dependent on cell death

The action of drugs in cell metabolism-based screening tests may depend on partial growth inhibition and cytostasis, besides cytotoxicity (23). To determine if the synergistic action of the SN38/erlotinib combination is due to increased cell death or decreased proliferation rate, we used the FLICK assay (Figure 2A) (24). The degree of cell death was evaluated using the Fractional Viability (FV) metric, which is simply the fraction of the total live cell population. FV analysis showed that the high potency achieved by the SN38/erlotinib combination was due to enhanced cell death (Figure 2B, Figure S2A). To determine if the levels of death observed were sufficient to shrink a tumor population, we also evaluated the normalized growth rate inhibition value (GR value). Negative GR values revealed that the SN38/erlotinib combination induced a significant shrinkage in population size, indicating high cell death (Figure 2C). In contrast, as indicated by positive GR values, erlotinib alone caused only a partial growth arrest, whereas SN38 led to partial growth arrest at low doses and cell death at high doses. We also evaluated the lethal fraction (LF) over time using FLICK to gain further insight into the kinetics of the death rate induced by these drugs. The SN38/erlotinib combination triggered cell death more robustly than single-agent treatments (Figure 2D-F, Figure S2B-C). CI analysis based on the LF metric revealed a strong synergism (Figure 2G), similar to CI values inferred from the RV-based analysis (Figure 1E). Hence, the LF-based assessment further validated that synergism in the SN38/erlotinib combination relied on an increased cell death rate.

**Figure 2.**
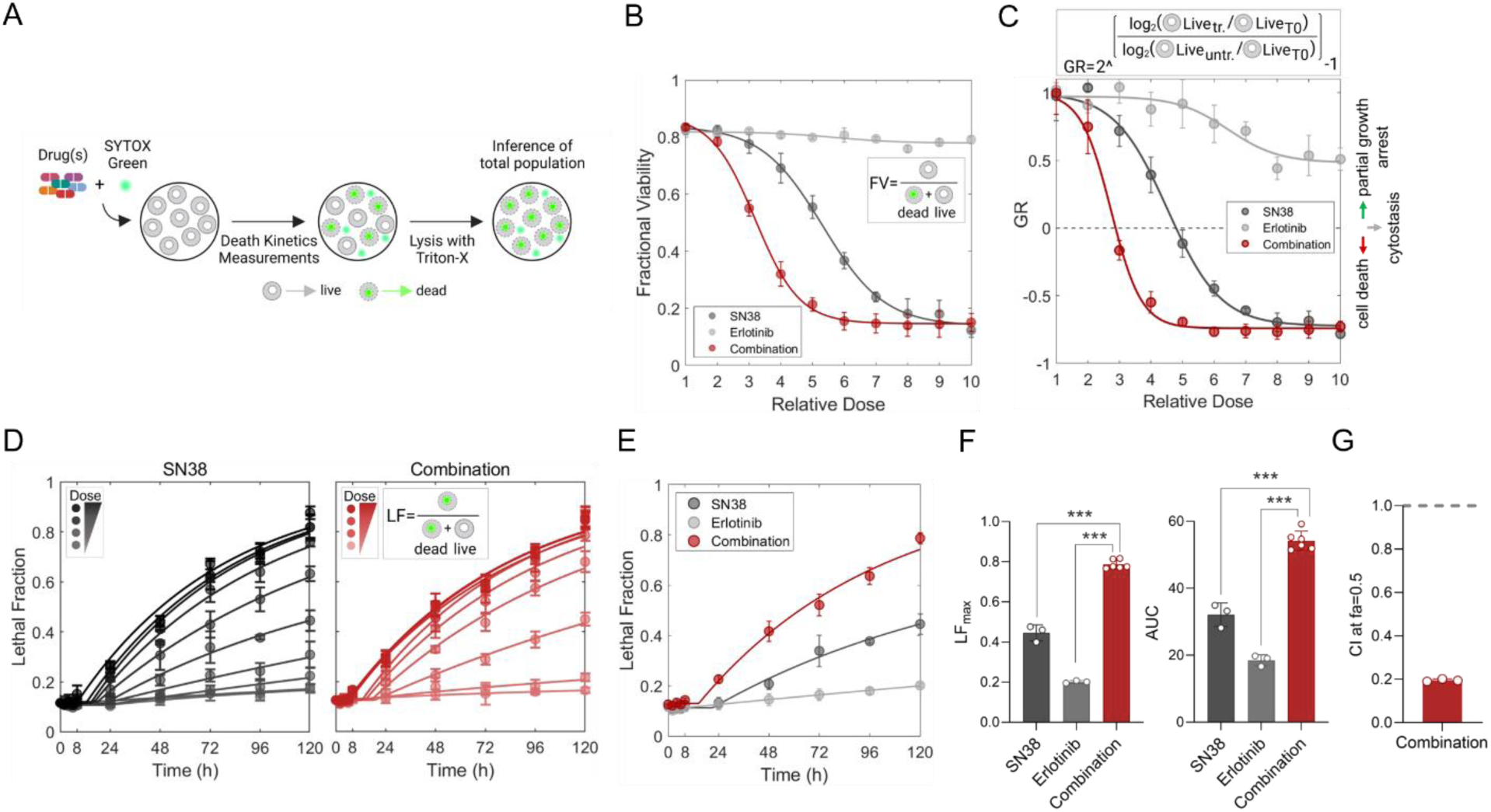
Validating the synergistic action of erlotinib-SN38 with the fluorescence-based and lysis-dependent inference of cell death kinetics (FLICK) assay. **(A)** The workflow for the FLICK assay adapted from Honeywell et. al. (25), created in BioRender. (**B)** Fractional viability (FV), **(C)** growth rate (GR), and **(D)** lethal fraction (LF) kinetics of the SN38/erlotinib combination and single-agent treatments. (B-D) Relative doses are given in Figure S2A. (C) Negative GR values indicate cell death and positive GR values indicate partial growth arrest. Untr: untreated, tr: treated. **(E)** Comparison of the LF curves for each treatment at relative dose 5. LF curves at all the relative doses are presented in Figure S2C. **(F)** Quantifying LF maxima (LFmax) and area under the curve (AUC) values of data presented in E. Statistical analysis was performed using Dunnett’s multiple comparisons test. *** *p*<0.001. **(G)** CI at fa:0.5, calculated based on LF metric.

To investigate growth arrest in detail, we analyzed the cell cycle progression under SN38 alone or SN38/erlotinib combination at relatively low and high doses. The low-dose SN38/erlotinib combination elicited a cell cycle arrest in S-phase, also observed for high-dose SN38 alone (Figure 3A, Figure S3A). However, the high-dose SN38/erlotinib combination induced a potent cell cycle arrest in the G1 phase and increased DNA damage even at earlier time points (Figure 3A-B, Figure S3A-B). Next, we explored the mode of synergistic cell death by investigating the alteration in cell death kinetics in the presence of extrinsic apoptosis, apoptosis, ferroptosis, necrosis, or parthanatos inhibitors (Figure 3C). Apoptosis inhibitor Z-VAD-FMK substantially decreased the LF under both SN38 and the SN38/erlotinib combination treatments, with a significantly delayed death onset (Figure 3C-D). The incapability of the extrinsic apoptosis-specific inhibitor Z-IETD-FMK to induce a similar change indicated that the cell death by the SN38/erlotinib combination was through intrinsic apoptosis. The assessment of cleaved-caspase3 and cleaved-PARP further confirmed prominent apoptotic cell death (Figure 3E, Figure S3C).

**Figure 3.**
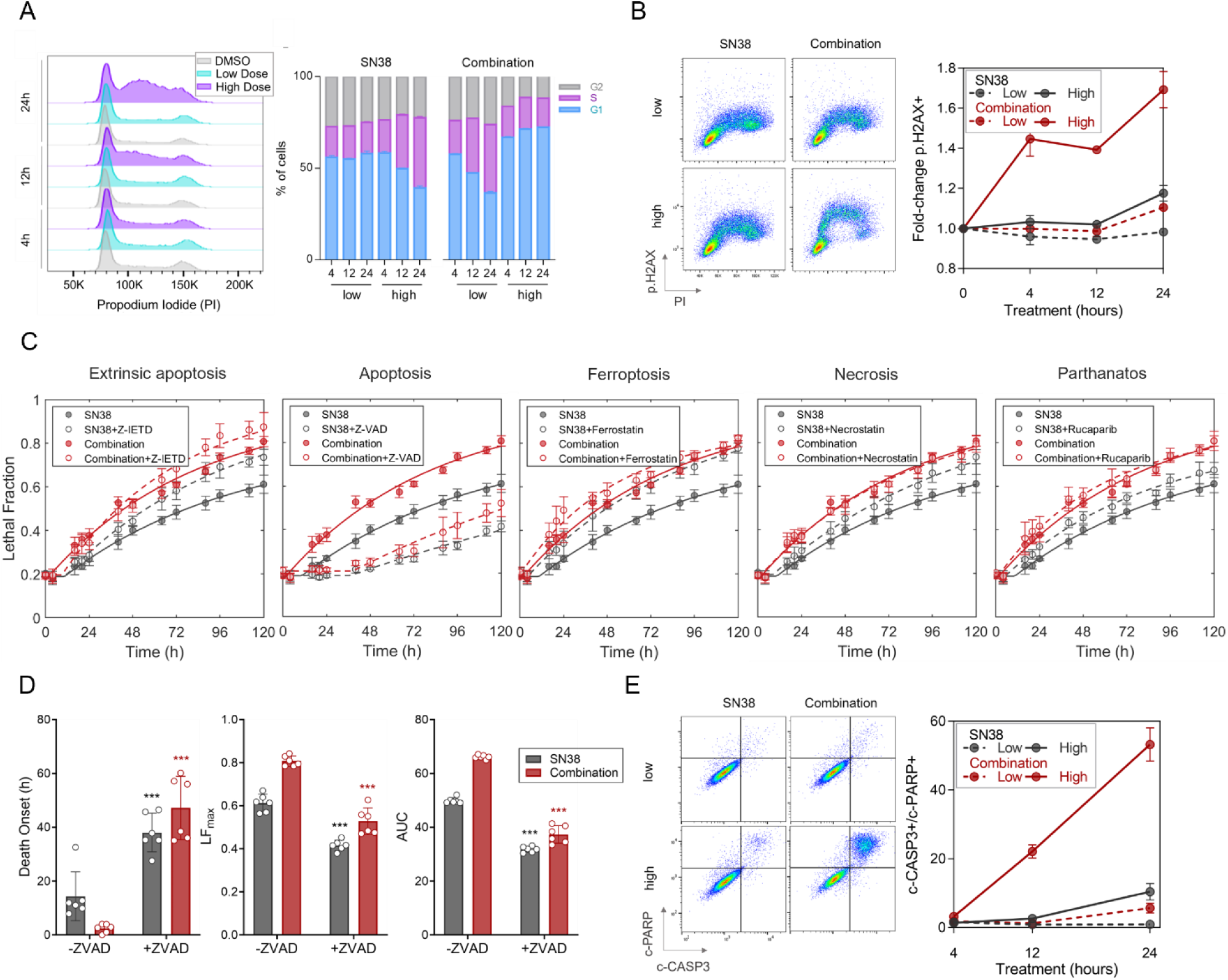
Phenotypic responses to SN38 alone and SN38/erlotinib combination in SNU5 cells. **(A)** Representative histogram for cell cycle progression (left), and percentage of cells at each phase (right), **(B)** representative flow cytometry plots (left) and fold-change in p.H2AX (right) to assess alterations in DNA damage over time under SN38 or SN38/erlotinib combination treatment at low or high doses. PI: propidium iodide. p.H2AX: phospho-H2AX. **(C)** Identification of cell death type triggered by SN38 alone (30nM) and SN38/erlotinib combination (30nM SN38 + 3μM erlotinib). Lethal fraction (LF) kinetics were analyzed over time under both treatments in the presence and absence of indicated inhibitors targeting specified cell death pathways. **(D)** The death onset time, LF maxima, and AUC parameters of LF curves reporting changes in cell death rate in the presence of apoptosis inhibitor Z-VAD-FMK in C. Statistical analysis was performed using Welch’s t-test. *** *p*<0.001. **(E)** Analysis of apoptotic response to SN38 alone or SN38/erlotinib combination treatment at low and high doses via cleaved-caspase3 (c-CASP3) and cleaved-PARP (c-PARP) co-staining. Representative flow cytometry plots (left) and the kinetics of apoptotic cell death quantified (right).

### The role of EGFR in synergistic action

To investigate the dependency of synergism on the inhibition of EGFR, the primary target of erlotinib, we analyzed the fractional viability of SNU5^EGFR-KO^ cells in response to SN38, erlotinib, and SN38/erlotinib combination (Figure 4A, B), and the kinetics of cell death via LF metric (Figure 4C). Unexpectedly, we did not observe a reversal of the synergism elicited by the SN38/erlotinib combination in SNU5^EGFR-KO^ cells compared to SNU5 parental cells, as FV in SNU5^EGFR-KO^ cells was indifferent from SNU5 cells under each treatment condition (Figure 4A-B). The evaluation of the LF also revealed that the SN38/erlotinib combination elicited a similar cell death rate and kinetics in both SNU5^EGFR-KO^ and SNU5 cells (Figure 4C). These results suggested a mechanism of synergism independent of EGFR inhibition.

**Figure 4.**
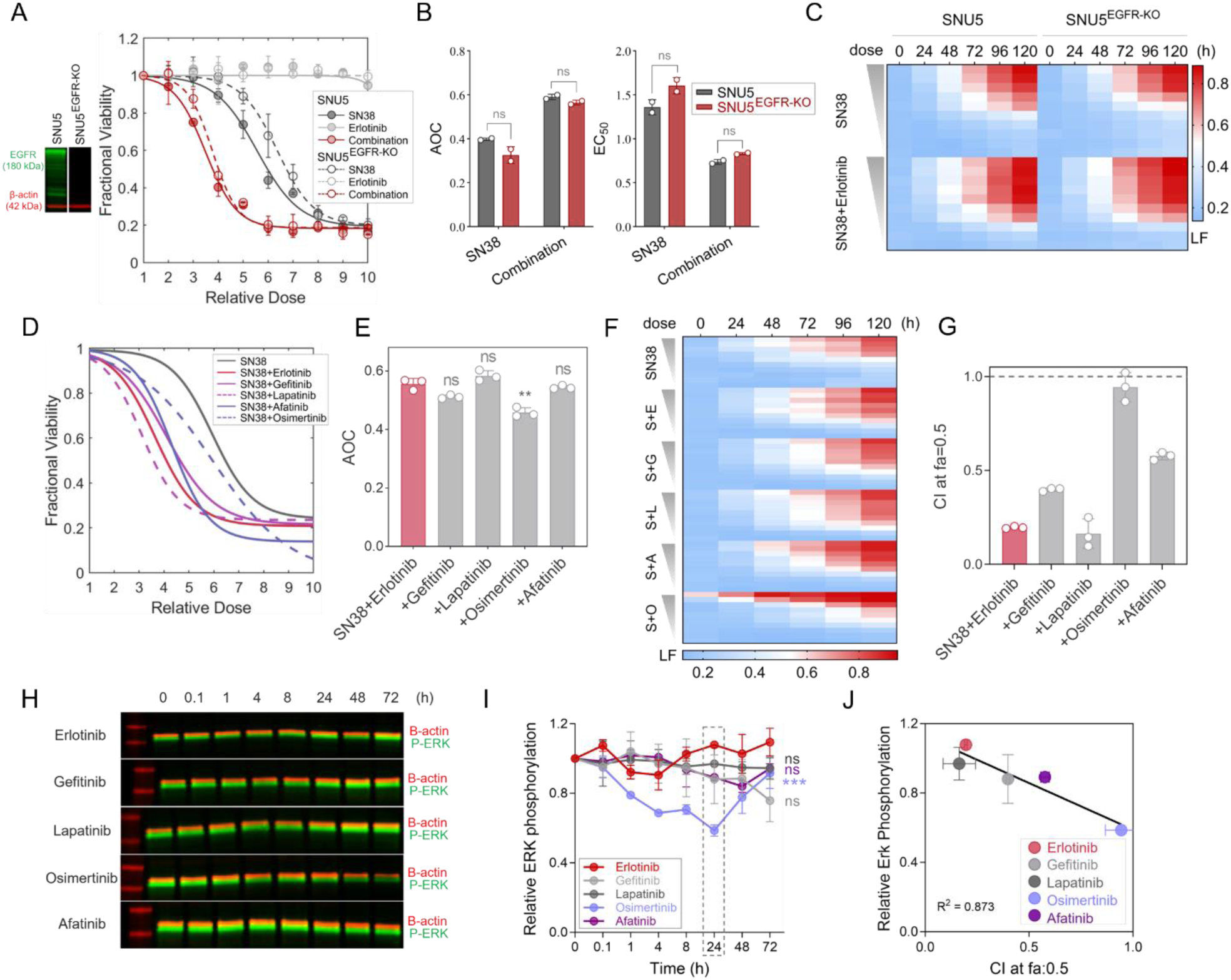
EGFR is not involved in synergistic action. **(A)** Drug response to the SN38/erlotinib combination and single agents in SNU^EGFR-KO^ cells compared to SNU5 cells. Immunoblot (left) shows the lack of EGFR expression in SNU^EGFR-KO^ cells. **(B)** Comparison of the area over the curve (AOC) and EC50 parameters deduced from fractional viability (FV) curves in A. Welch’s t-test was used for statistical analysis. ns: non-significant. **(C)** Dose-dependent cell death kinetics over time in SNU^EGFR-KO^ and SNU5 cells. **(D)** Drug response to SN38 and its dual combination with EGFR inhibitors. **(E)** Comparison of the AOC values deduced from FV curves in D, using Dunnett’s multiple comparisons test. ** p-value=0.002. **(F)** Dose-dependent cell death kinetics over time under SN38 alone and its dual combination with EGFR inhibitors. (S: SN38, E: Erlotinib, G: Gefitinib, L: Lapatinib, O: Osimertinib, A: Afatinib). **(G)** CI values at fa:0.5 deduced from LF analysis in F. **(H)** Immunoblot analysis of time-dependent alterations in ERK activity under treatment with EGFR inhibitors. **(I)** Densitometric quantitation of immunoblot data in H. The dashed box indicates the time point where a significant decrease in ERK phosphorylation was observed under osimertinib treatment, but not other EGFR inhibitors, in comparison with erlotinib. Statistical significance was determined using Dunnett’s multiple comparisons test. *** p<0.001. **(J)** Correlation between the level of ERK phosphorylation at 24 hours after treatment with the specified EGFR inhibitor and CI value at fa:0.5 for dual combinations of SN38 with EGFR inhibitors.

Several studies report different efficacies and toxicity profiles for distinct EGFR inhibitors in cancer patients (26). To understand whether the synergistic interaction of erlotinib with SN38 is common to different EGFR inhibitors, we tested the combinations of gefitinib, lapatinib, osimertinib, and afatinib with SN38 in SNU5 cells (Figure 4D-F, Figure S4A). The dual combination of SN38 with all EGFR inhibitors improved the drug response and enhanced the cell death rate compared to SN38 alone, as in the SN38/erlotinib combination, except for osimertinib. CI analysis showed that all EGFR inhibitors, but not osimertinib, exhibited synergism with SN38 (Figure 4G). To assess whether EGFR inhibitors’ action on other receptor tyrosine kinases (RTK) may be involved in synergism, we examined the kinetics of ERK phosphorylation, a common intracellular target of the RTK family (27) (Figure 4H-I, Figure S4B). Except for Osimertinib, the EGFR inhibitors did not decrease ERK phosphorylation in SNU5 cells, ruling out the involvement of other RTKs as off-targets of EGFR inhibitors in synergistic response. Surprisingly, the degree of synergism with SN38 was inversely correlated with the efficacy of EGFR inhibitors to inhibit ERK phosphorylation (Figure 4J). Since EGFR or RTK signaling through pERK is not altered by EGFR inhibitors that elicited synergism with SN38, these findings strengthened that the synergism likely does not result from the loss of EGFR or RTK signaling.

### Investigating the genetic dependency/vulnerability signature for drug synergy with whole-genome CRISPR screening

To elucidate the distinct genetic dependencies underlying cell death activation via the SN38/erlotinib combination compared to SN38 alone, we adopted a comprehensive approach, integrating genome-wide CRISPR screening with annexin-V magnetic bead sorting (Figure 5A). As shown in Figures 2E and 3E, SN38/erlotinib combination induces cell death more potently than SN38 alone when we apply SN38 at equimolar concentrations in both treatments. However, in equimolar concentrations, it is hard to discern whether the SN38/erlotinib combination employs distinct mechanisms of action than SN38 alone or if erlotinib’s presence reinforces the exact mechanisms of action triggered by SN38 for cell death. Therefore, we identified equipotent concentrations of SN38 and SN38/erlotinib combination that achieve an intermediate level of cell death at a shorter drug exposure period, as opposed to previous CRISPR-based perturbation screens with DNA-damaging agents (28,29). Such adjustments ensured that the population size that would guarantee sgRNA representation at >300 coverage could be maintained throughout the screen, and the sgRNA distribution of the library would mainly be affected by the alterations in cell death rate, instead of the alterations in proliferation rate. Exposure to 13.5nM SN38 and the combination of 3.7nM SN38 and 370nM erlotinib for three days induced ̴ %50 annexin-V positivity (Figure 5B-D). We applied these assay conditions in our pooled screen to unveil the mechanistic traits intrinsic to cell death triggered by SN38 and SN38/erlotinib combination.

**Figure 5.**
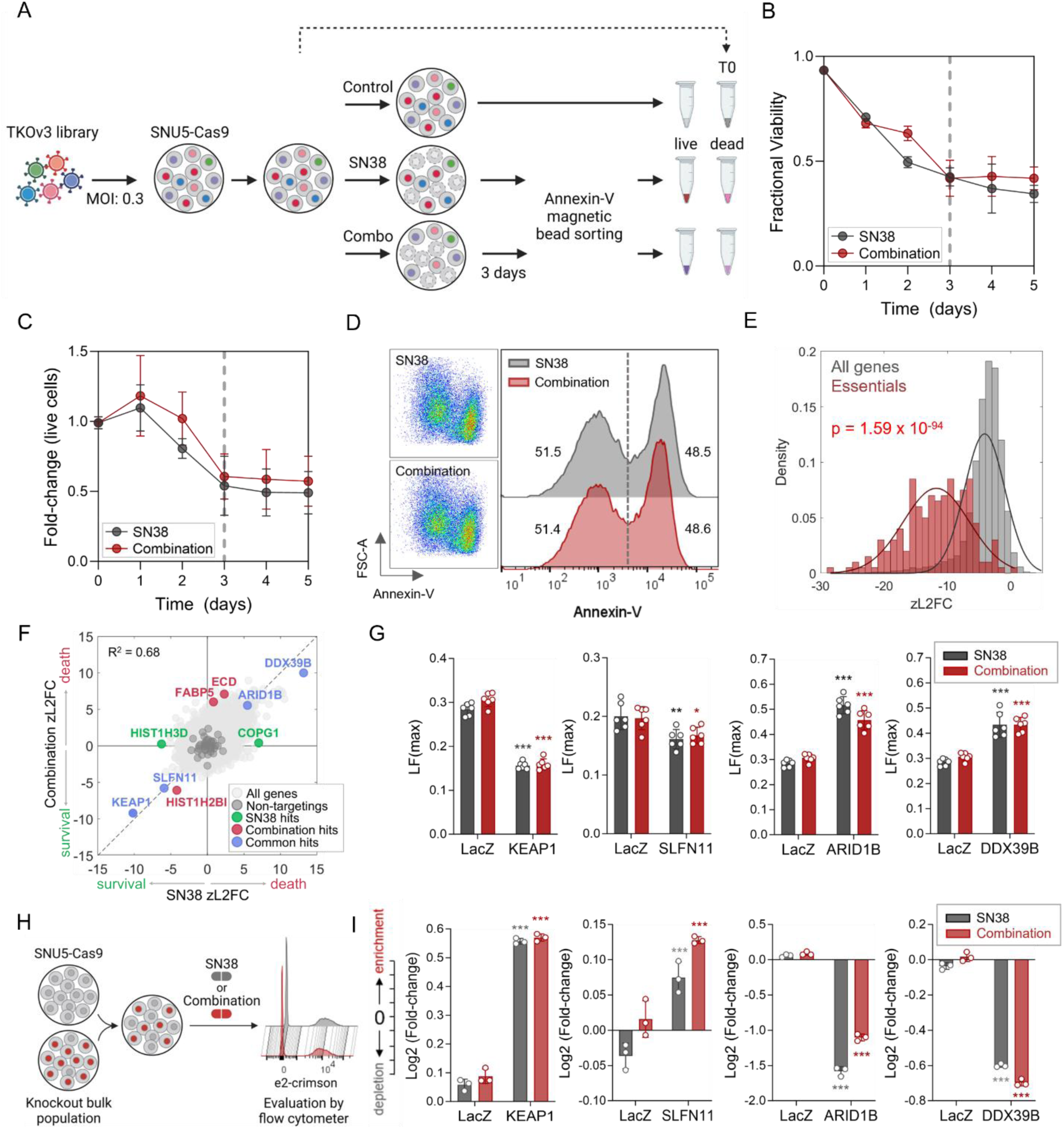
Genome-wide CRISPR screen to explore the mechanism of synergism. **(A)** Experimental workflow of the genome-wide CRISPR screening coupled with annexin-V magnetic bead sorting. The figure was generated using BioRender. **(B-C)** Optimization of SN38 (13.5nM) and the SN38/erlotinib combination (3.7nM SN38+370nM erlotinib) concentrations and drug exposure period for CRISPR screen based on trypan-blue exclusion. Treatment with SN38 alone and the SN38/erlotinib at indicated concentrations for 3 days resulted in (B) fractional viability of ̴ 0.5 obtained by counting both the number of live and dead cells and (C) a ̴ 0.5-fold change in the number of live cells in the treated group compared to the untreated. **(D)** Evaluation of annexin-V positivity under SN38 alone and the SN38/erlotinib combination applied at optimized concentrations for 3 days. The assay conditions for both treatments achieved ̴ %50 annexin positivity. **(E)** Distribution of all genes compared to core essential genes in untreated vs. T0 sample at the gene-level zL2FC. The Kolmogorov-Smirnov test was used to calculate the p-value. **(F)** Identification of candidate hit genes with an FDR<0.1 for both SN38 alone and the SN38/erlotinib combination treatments, comparing the dead vs. the live populations at the gene-level zL2FC. The knockout of genes highlighted significantly altered cell death or survival rates under SN38 alone and/or the SN38/erlotinib. Positive zL2FC values indicate an increase in death rate, and negative zL2FC values indicate an increase in survival rate. The relationship between the two data sets was determined by computing R-squared for the genes with an FDR<0.1. **(G)** Investigating alterations in the death rate within the knockout populations compared to the untargeted population (LacZ) under SN38 and SN38/erlotinib combination by analyzing LFmax. **(H)** The schematic representation of the competition assay generated using BioRender. **(I)** Results of competition assay showing the enrichment/depletion of the knockout populations compared to the untargeted population under SN38 and SN38/erlotinib combination. The treatments were applied at the screening doses for 3 days in (G) and (I). Dunnett’s multiple comparisons test was used to compare the knockout and untargeted populations for each treatment. *** p<0.001, ** 0.002, * 0.033.

The quality assessment of the screen data revealed a significant decrease in core essential genes (30) in the untreated vs. T0 sample and a strong correlation between replicates (Figure 5E, Figure S5A). We followed a CRISPR death screen approach (31), using the comparison of “dead vs. live” at the gene-level z-scored log_2_ fold change (zL2FC) for the SN38/erlotinib combination compared to SN38 to identify differentially enriched or depleted genes under either of these treatments and both (Figure 5F). The gene-level knockouts significantly enriched/depleted with an FDR<0.1 in the SN38/erlotinib combination showed a strong correlation with those in SN38 alone. The evaluation of the LF metric under SN38 alone and the SN38/erlotinib combination demonstrated that deleting KEAP1 or SLFN11 decreased the death rate. In contrast, deleting ARID1B or DDX39B enhanced the death rate compared to the untargeted population, validating the screen results (Figure 5G). We performed competition assay in knockout populations to provide insight into the altered cell survival in response to treatments. We observed the enrichment of KEAP1 or SLFN11 knockout populations, as opposed to the depletion of ARID1B and DDX39B knockout populations in a mixed culture of SNU5-Cas9 and knockout populations under both treatments (Figure 5H-I). However, the knockout of the genes that altered cell death or survival rates under either SN38 or the SN38/erlotinib combination, but not both, failed to validate the screen data (Figure S5C-D). The high correlation between experiment groups treated with SN38 alone and the SN38/erlotinib combination and validation experiments suggested remarkably similar genetic dependency/vulnerability signatures for these treatments, indicating shared mechanisms of action.

### Dissecting the mechanism of synergism with RNAi-based signature assay

To validate whether the SN38/erlotinib combination employs the exact mechanisms of action with SN38 alone, we conducted an RNAi-based signature assay, as it provides both statistical and biological generalization for the actions of anti-cancer agents (5,7,20). For this, we generated the signatures of the SN38/erlotinib combination and each single-agent treatment by assembling resistance index (RI) values for each cell population expressing one of the eight shRNAs. Then, we compared the signatures to a reference set using the modified K-nearest neighbors algorithm (Figure 6A, Figure S6A). Our results demonstrated that the SN38/erlotinib combination, like SN38, clustered in the TOP1 poison category, with a linkage ratio of 1 and p-value <0.05 (Figure 6B, Figure S6B-C). Thus, this analysis further confirmed that the SN38/erlotinib combination operated through the exact mechanisms of action of SN38, as indicated by both biological and statistical classifications.

**Figure 6.**
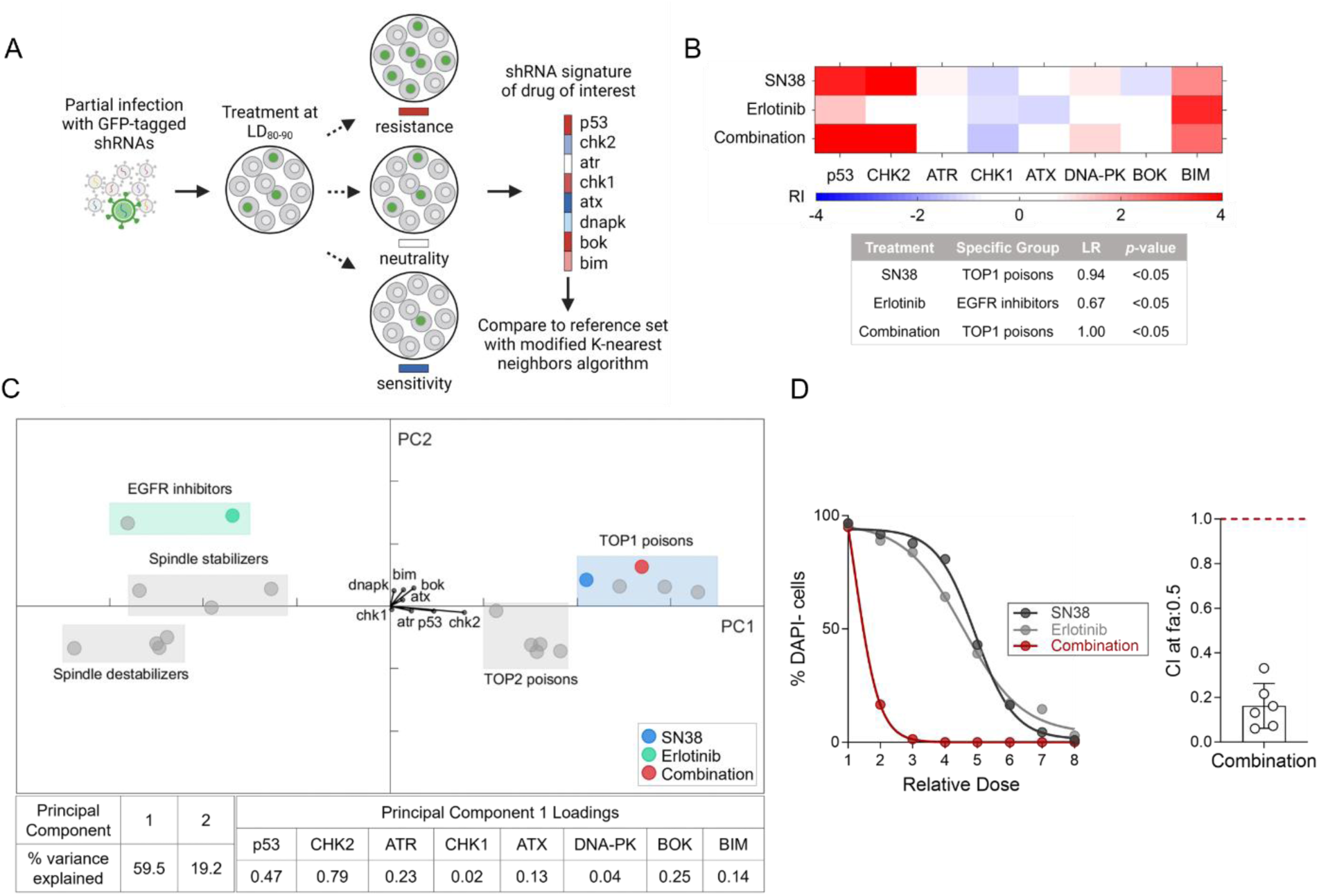
Investigating the mechanism of synergism with the shRNA-based signature assay. **(A)** Illustration of RNAi-based signature assay, adapted from Jiang et al. (7), using BioRender. LD80-90: lethal dose inducing 80-90% cell death. **(B)** The heatmap generated by assembling resistance index (RI) values for each cell population expressing the specified shRNA shows the signatures of the single agents and SN38/erlotinib combination (top). Each drug signature was compared with the reference set using the modified K-nearest neighbors algorithm that reports the linkage ratio (LR) and p-value (bottom). **(C)** Principal component analysis of TOP1 and TOP2A poisons, EGFR inhibitors, spindle stabilizers, and destabilizers. Boxes in the graph highlight the estimated spatial position of each category within the PCA. The percent variance explained by each principal component and principal component 1 loadings showing each shRNA’s contribution to the analysis is provided in the table. **(D)** Drug response analysis of the SN38/erlotinib combination and single agents in Eμ-Myc Cdkn2a^Arf−/−^ cells (left), and the calculation of CI at fa:0.5 for the SN38/erlotinib combination (right).

We performed principal component analysis to examine the variance in our data in fewer dimensions (Figure 6C), showing the separation of EGFR inhibitors and TOP1 poisons along the first principal component (PC1). Among the original variables from which PC1 was established, shCHK2 strongly contributed to PC1 compared to other hairpins. CHK2 is one of the crucial regulators of the signaling cascade that conveys the DNA damage signal to various downstream effectors (32). Therefore, this data indicated that the SN38/erlotinib combination induced DNA damage response, like other members of the TOP1 and TOP2A poisons categories, supporting our previous data (Figure 3B). We also confirmed a strong synergism between SN38 and erlotinib in Eμ-Myc Cdkn2a^Arf−/−^ leukemia cells, used as a cell model in the signature assay (Figure 6D, Figure S6E-F).

### Exploring the off-target effects of Erlotinib

Given that EGFR or other RTKs are not involved in the synergistic interaction of erlotinib with SN38, and the SN38/erlotinib combination exhibited the same genetic dependency signature as SN38, we reviewed the literature for other targets of EGFR inhibitors that can potentiate the action of SN38. Since previous studies reported several efflux pumps as off-targets of EGFR inhibitors (33), we reassessed our genome-wide screening data to find whether the knockout of any efflux pump altered the cell death or survival rate under SN38 only and combination treatments. We observed that the genetic ablation of ABCG2, also known as breast cancer resistance protein (BCRP) (34), enhanced the cell death rate in the screen under both treatments (Figure 7A).

**Figure 7.**
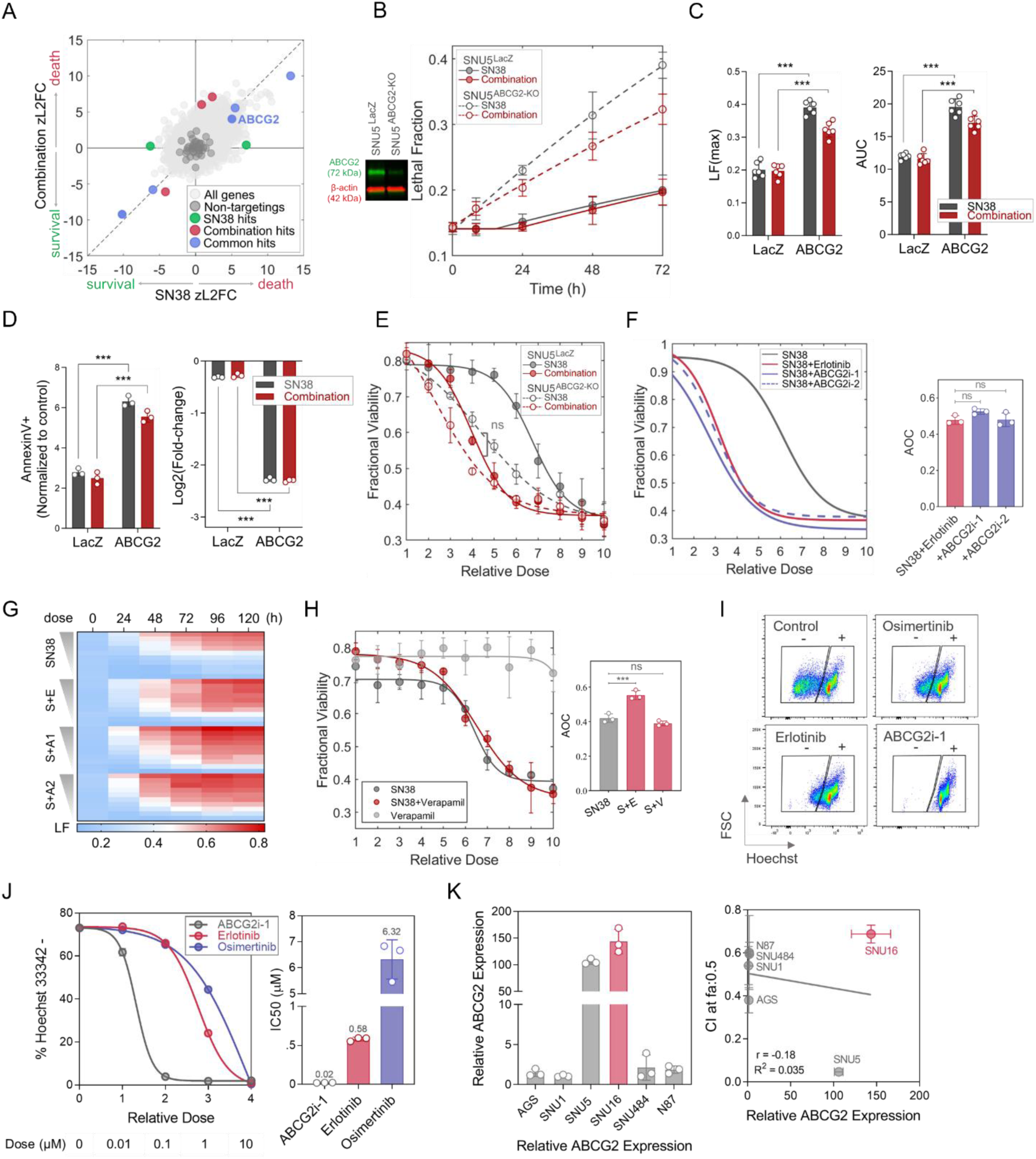
The erlotinib’s off-target effect on ABCG2 is responsible for synergism with SN38. **(A)** The screen result shows an enhanced cell death rate under SN38 and the SN38/erlotinib combination with the knockout of ABCG2. **(B)** Analysis of cell death kinetics via lethal fraction (LF) and **(C)** LFmax plots derived from (B). Immunoblot in B (left) shows decreased ABCG2 expression in the SNU5^ABCG2-KO^ bulk population compared to SNU5^LacZ^. **(D)** Annexin V positivity (left) and survival ability assessed by competition assay (right) in SNU5^ABCG2-KO^ compared to SNU5^LacZ^ cells under SN38 alone and the SN38/erlotinib combination. **(E)** Evaluating drug response to SN38 alone and the SN38/erlotinib combination in SNU5^ABCG2-KO^ and SNU5^LacZ^ cells using fractional viability (FV). **(F-G)** Dual combination of SN38 with ABGC2 inhibitors (ABCG2i-1 and ABCG2i-2) or with erlotinib. (F) Drug response analysis using FV metric (left) and the comparison of AOC values deduced from FV curves (right) in SNU5 cells. (G) Dose-dependent cell death kinetics over time. A1: ABCG2i-1, A2: ABCG2i-2. **(H)** FV in response to the SN38/verapamil combination (left), comparison of AOC values (right). S: SN38, E: erlotinib, V: verapamil. **(I-J)** Assessment of intracellular Hoechst-33342 accumulation in a dose- and time-dependent manner under erlotinib, osimertinib, or ABCG2i-1 treatments compared to the control. (I) Representative flow plots (1h). (J) Dose-dependent Hoechst-33342 negativity at the end of the 16-hour efflux period (left) and IC50 values (right). **(K)** Relative ABCG2 expression in six gastric cancer cell lines (left) and its correlation with CI at fa:0.5 values for the SN38/erlotinib combination (right). Welch’s t-test was used for all the panels with statistical testing. ***p-value<0.001. ns: non-significant.

By adopting different validation approaches, we demonstrated that knocking out the ABCG2 gene enhanced the drug-induced cell death and impaired the survival ability under SN38 and the SN38/erlotinib combination, which confirmed the screen finding (Figure 7B-D, Figure S7A). Moreover, SN38-only treatment in SNU5^ABCG2-KO^ cells phenocopied the drug response to the SN38/erlotinib combination in SNU5^LacZ^ cells (Figure 7E). Next, we investigated the drug response to the dual combination of SN38 with different ABCG2 inhibitors, ABCG2i-1 (KO143) and ABCG2i-2 (KS176) in SNU5 cells. The combination of SN38 with either ABCG2i-1 or ABCG2i-2 (Figure 7F-G, Figure S7B-C) achieved a similar degree of drug response and cell death to the SN38/erlotinib combination. However, the ABCB1 inhibitor verapamil could not increase the response to SN38 (Figure 7H). Hence, these results suggested that the synergism in the SN38/erlotinib combination exhibited a specific dependence on the inhibition of ABCG2 efflux pump activity by erlotinib.

We then investigated the effect of erlotinib on the ABCG2 efflux pump activity in a dose- and time-dependent manner by analyzing the intracellular accumulation of Hoechst-33342 dye, a substrate for ABCG2 (Figure 7I-J, Figure S7D-E). We used ABCG2i-1 as the positive control. Since the drug response and immunoblot findings suggested osimertinib as the only EGFR inhibitor with action on RTK activity and no synergism with SN38 in SNU5 cells, we employed osimertinib as a negative control treatment (Figure 4D-I). Erlotinib inhibited the ABCG2 efflux activity, but its effect was not as strong as ABCG2i-1, as expected (Figure 7I-J, Figure S7D-E). Osimertinib also had an inhibitory effect on the efflux pump activity at high concentrations. However, nearly an 11-fold higher dose of osimertinib was required to achieve the same degree of ABCG2 inhibition as erlotinib. These findings proved that erlotinib was a potent inhibitor of the ABCG2 efflux pump in SNU5 cells, explaining why the SN38/osimertinib pair failed to induce a strong synergism compared to the SN38/erlotinib combination. Moreover, the degree of synergism positively correlated with the ABCG2 expression level for gastric adenocarcinoma cell lines, except for SNU16 cells (Figure 7K). These findings suggested that the expression level of ABCG2 could serve as one of the biomarkers for predicting sensitivity to the SN38/erlotinib combination.

## Discussion

The mainstay of systemic therapy for cancer is the combination of anticancer drugs with non-overlapping mechanisms and toxicities to increase anticancer efficacy, decrease toxicity, address tumor heterogeneity, and prevent chemoresistance (3). Achieving these goals requires a comprehensive understanding of mechanisms of action for single drugs/combinations, and dissecting the genetic dependencies for pharmacological interactions (4). Although the basis for such knowledge is being established for the combinations of conventional chemotherapeutics (5,6), the mechanisms by which molecular-targeted agents interact with chemotherapeutics are yet to be discovered. Hence, in this study, we performed a pairwise screening of molecular-targeted agents with conventional chemotherapeutics in gastric adenocarcinoma cells since gastric cancer is an intractable cancer with an urgent need for a rational design of combination therapies.

Our pairwise pharmacological screen identified the SN38/erlotinib as a drug pair with a robust synergism in gastric adenocarcinoma cells. The synergistic interaction between the two was much stronger than the currently available combination regimens in gastric cancer, consistent across different assays, and relied on an enhanced cell death rate. Unexpectedly, however, the synergism between SN38 and erlotinib did not depend on the primary target of erlotinib, EGFR, directing us to perform a genome-wide CRISPR screening and shRNA-based signature assay to decipher the underlying genetic signature of the synergism. Such an approach was also crucial to identify more robust targets that low doses of available agents can inhibit. Because a lower potency of erlotinib compared to SN38, which requires high doses to achieve synergism, can be a limitation for use in the clinic. Our functional genomics approaches integrated with multistep validation assays showed that the genetic dependency signature of the SN38/erlotinib combination was the same as SN38. Although this result may be expected considering the independence of synergism from EGFR, it was still surprising since EGFR inhibitors inhibit several other RTKs in cancer cells (35), which would reprogram intracellular machinery, leading to distinctive dependency signatures.

To identify the critical player for synergism in the SN38/erlotinib combination, we revisited our screen data for genes, targeting which may enhance the action of SN38 without changing the genetic dependency signature. Pharmacological knowledge of cancer chemoresistance indicates that drug efflux pumps can serve such functions. Accordingly, emerging evidence supports that the ABC transporter superfamily can be off-targets of EGFR inhibitors in cancer (33). Among the ABC transporters, the ABCG2 emerged as a hit enriched in the dead population in both SN38 monotherapy and SN38/erlotinib combination groups in our CRISPR screen. Gene knockout and pharmacological inhibition studies also confirmed the inhibition of ABCG2 as an off-target of erlotinib as the mechanism of synergism with SN38 in SNU5, derived from a gastric adenocarcinoma patient who had previously received chemotherapy.

Since ABCG2 is involved in the efflux of a broad range of chemotherapeutics, translating ABCG2 inhibitors into the clinic is an attractive strategy in cancer (34). While some ABCG2 inhibitors have been tested in clinical trials (36–38), tyrosine kinase inhibitors are also emerging as ABCG2 inhibitors (33). Therefore, our findings may support the use of ABCG2 inhibitors or EGFR inhibitors in combination with SN38 or irinotecan in gastric adenocarcinoma. This strategy may achieve a significant tumor size reduction in ABCG2-positive tumors in the first-line setting. Moreover, in the second-line setting, it may be efficacious to kill clones that become resistant after first-line chemotherapy due to the upregulation of ABCG2. EGFR inhibitors may also provide an advantage in acting both on cells susceptible to the action of erlotinib on EGFR and cells sensitive to the off-target activity on ABCG2 in highly heterogeneous gastric adenocarcinomas.

Our study firmly demonstrated how functional genomics approaches can uncover biologically relevant and validated mechanisms underlying drug action and synergistic interactions in combination therapy. Even though we used different functional genomics approaches that utilize different modes of genetic ablation and a substantially different extent of targeted genes (7,39), both the genome-wide screening and the shRNA-based signature assay indicated that the genetic signature of the SN38/erlotinib combination is not different from the single agent SN38. In addition, we used the FLICK assay (15) and competition assay as powerful tools to validate the synergistic interaction in cell death and the functional significance of the hit genes in the genetic signature. Thus, we reliably inferred the mechanism of synergism of a promising drug pair in gastric adenocarcinoma.

The accurate quantitation of synergism among a set of drugs is only feasible in cell models. However, drugs may interact at multiple levels in animal models and patients, altering the overall efficacy-safety profile of combination regimens in the clinic. Therefore, using in vitro cell lines to investigate drug synergy in this study may be a limitation. For instance, the SN38/erlotinib combination may exhibit an increased risk of adverse effects since erlotinib and SN38 also interact pharmacokinetically at the metabolism level, increasing plasma concentrations of SN38 (40). This may increase the risk of dose-limiting toxicities of SN38. ABCG2 inhibitors may be more prone to a similar failure, requiring watchful dose-optimization studies. These points may also raise the question of whether molecular-targeted agent-conventional chemotherapeutic pairs that exhibit synergism via an averaged phenotype of genetic dependencies/vulnerabilities or generate a signature distinct from individual drugs would be more efficient and safer options in the clinic. This can be true for therapy response in the general patient population. However, inhibition of ABCG2 to enhance the action of SN38 can be a more effective strategy in tumors with high ABCG2 expression and sensitivity to the DNA-damaging action of SN38. These points should be addressed in future studies.

Other foci of our prospective studies will be on the selective efficacy of EGFR inhibitors on ABCG2 and alternative mechanisms for synergism. It is unclear why the EGFR inhibitors, except for osimertinib, fail to inhibit ERK phosphorylation in SNU5 cells despite these cells expressing wild-type EGFR. Besides, although the synergism was generally well correlated with the degree of ABCG2 expression, SNU16 cells with the highest ABCG2 expression but the lowest synergism disrupted this correlation. This data suggests that there can be additional dependencies for synergism, which is likely related to each cell line’s repertoire of mutational signatures. As supported by our perturbation screen and previous studies, decreased SLFN11 expression is responsible for drug resistance to DNA-damaging agents (41). Based on the publicly available data set (merav.wi.mit.edu/) (42), SLFN11 expression is significantly lower in SNU16 cells compared to SNU5 cells, which may be the cause of the breached correlation between ABCG2 expression and synergism (Figure S7F).

A clinically revealing part of this study is the insufficiency of the primary drug target to predict the synergism. Molecular-targeted agents are designed to target tumor-specific biomarkers with protumorigenic effect. Since the mutations in the target protein or the downstream signaling pathway are associated with the development of resistance to these agents, preclinical, translational, and precision medicine efforts are directed to catalog such mutations and use this knowledge for patient stratification (43). However, these approaches should advance with the evidence indicating off-targets as the main dependencies for the action of numerous molecular-targeted agents (9). Accordingly, our findings elucidate that proteins that are considered off-target can better predict response to combination therapy in cancers with specific genetic backgrounds. Therefore, a comprehensive understanding of the genetic dependency/vulnerability signatures and cell type-specific mutational landscape is crucial for a more accurate assessment of response to the molecular-targeted agent-conventional chemotherapeutic combinations. Our results support that the ABCG2 status should be assessed in addition to the EGFR status to improve patient selection strategies for EGFR inhibitor-conventional chemotherapeutic combinations.

In conclusion, our functional genomics approach elucidated that synergistic molecular-targeted agents can act by enhancing the action of conventional chemotherapeutics, leading to an identical genetic dependency/vulnerability profile in combination therapy as conventional chemotherapeutics. The emergence of an off-target, but not on-target, as the predictor of synergism implicates the significance of assessing the off-target status together with the on-target for a more precise selection of patient groups in clinical studies. Further dissection of the interaction between molecular-targeted agents and conventional chemotherapeutics may lead to the development of effective combination regimens for treating gastric adenocarcinoma, where the chances of cure are almost nil and anticancer agents are limited. Such efforts also have a high potential for discovering new targeted therapies that induce synergism with conventional chemotherapeutics.

### Author’s Disclosures

G. Ozcan reports that the pairwise screening of molecular-targeted agent-conventional chemotherapeutic pairs for synergism in gastric cancer cells is based on a former grant by The Scientific and Technological Research Council of Turkey (TUBITAK) with grant number 117Z460. M.J. Lee reports a grant from the National Institutes of Health/NIGMS (RO1 GM127559). M.E. Honeywell reports a grant from the NCI (F31 CA268847). M.T. Hemann reports grants from NCI (R01-CA233477) and NIH (R01-CA226898).

### Author’s Contributions

O.L. performed all the experiments and statistical analyses. O.L. and M.E.H. analyzed the CRISPR screening data. O.L. and G.O. wrote the manuscript. O.L., G.O., M.J.L., and M.T.H. conceptualized the research. G.O., M.J.L., and M.T.H. supervised the study. All authors reviewed and edited the manuscript.

## Acknowledgments

The authors gratefully acknowledge Hakan S. Orer and Mehmet Gonen from Koç University for their valuable insights as advisors in the project with grant number 117Z460-TUBITAK and the use of the services and facilities of the Koç University Research Center for Translational Medicine (KUTTAM), funded by the Presidency of Turkey, Head of Strategy and Budget. We also gratefully acknowledge Peter M. Bruno for his helpful comments on the RNAi-based signature assay, Tugba Bagci Onder for supportive insights, and Alisan Kayabolen, Tolga Sever, and James Ham for helpful discussions. We thank all members of Lee and Hemann Labs for their valuable feedback. O.L. gratefully acknowledges the support of Ph.D. Dissertation Research Grant sponsored by the U.S. Department of State and the Turkish Fulbright Commission. The contents presented in the manuscript are solely the responsibility of the authors and do not necessarily represent the official views of the Fulbright Program, the U.S. Department of State, and the Turkish Fulbright Commission.

## Notes

### Competing Interest Statement

The authors have declared no competing interest.

### Summary of Updates

This version of the manuscript has been revised to update the following: 1. The title was updated to clarify the message of the study. 2. Figure 2 and Figure 5 were updated to convey results more clearly. 3. The legends of Figures 2 and 6 were updated to cite references and include cross-references to supplementary figures. 4. The acknowledgement section was extended. 5. The legend of Figure S2 was updated to include cross-references to figures.

