## Supplementary Figures for "Functional genomics reveals an off-target dependency of drug synergy in gastric cancer therapy"

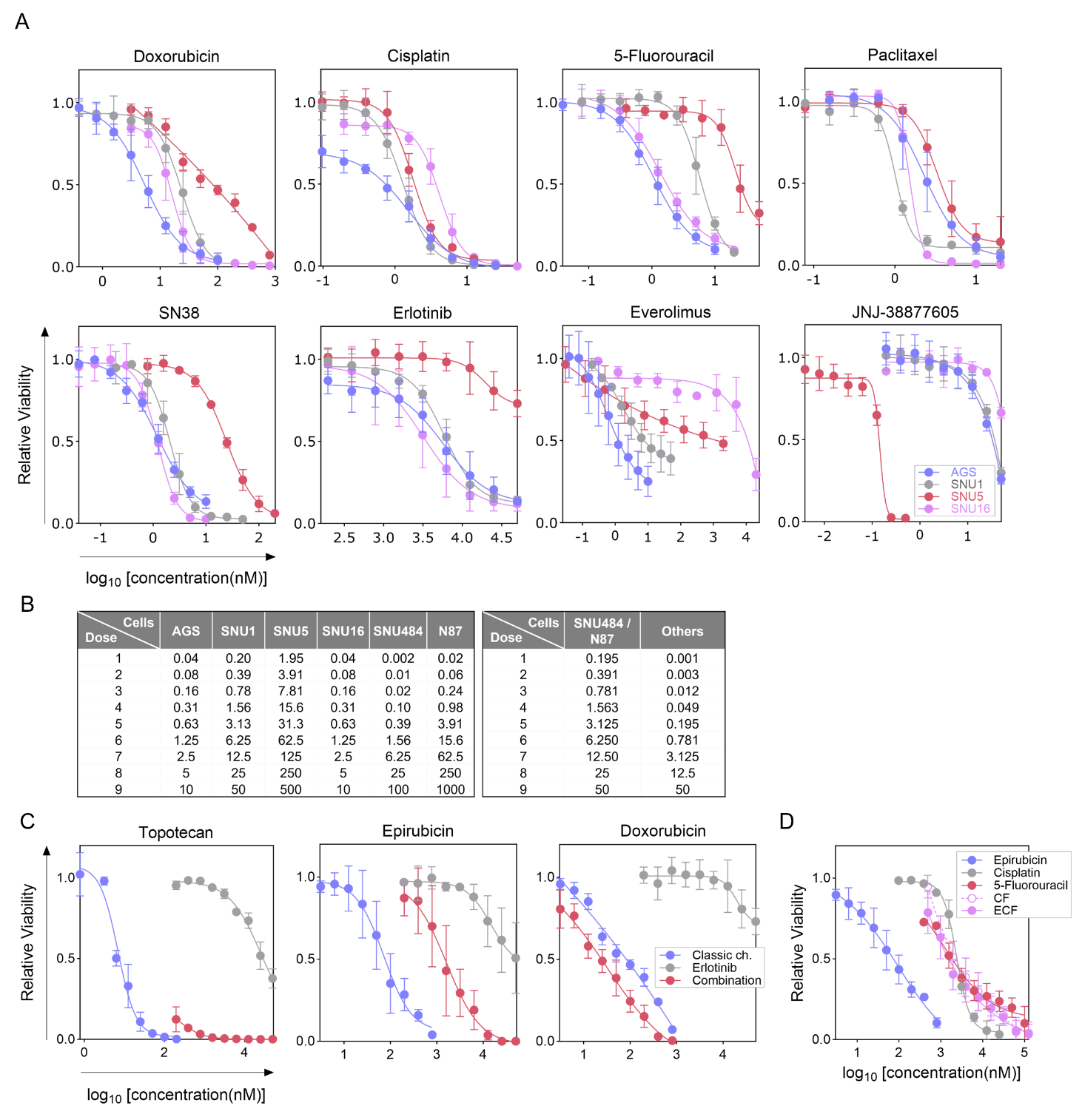

**Figure S1. (A)** The dose-response curves for conventional chemotherapeutics and molecular-targeted agents in gastric adenocarcinoma cells. **(B)** The relative doses for SN38 in nM (left) and erlotinib in µM (right) used in the combination experiments presented in Figure 1C. **(C)** The dose-relative viability curves for the combination of erlotinib with topotecan, epirubicin, and doxorubicin in SNU5 cells**. (D)** The dose-relative viability curves for epirubicin, cisplatin, 5-fluorouracil, CF (cisplatin and 5-fluorouracil combination), and ECF (epirubicin, cisplatin, 5-fluorouracil) regimen.

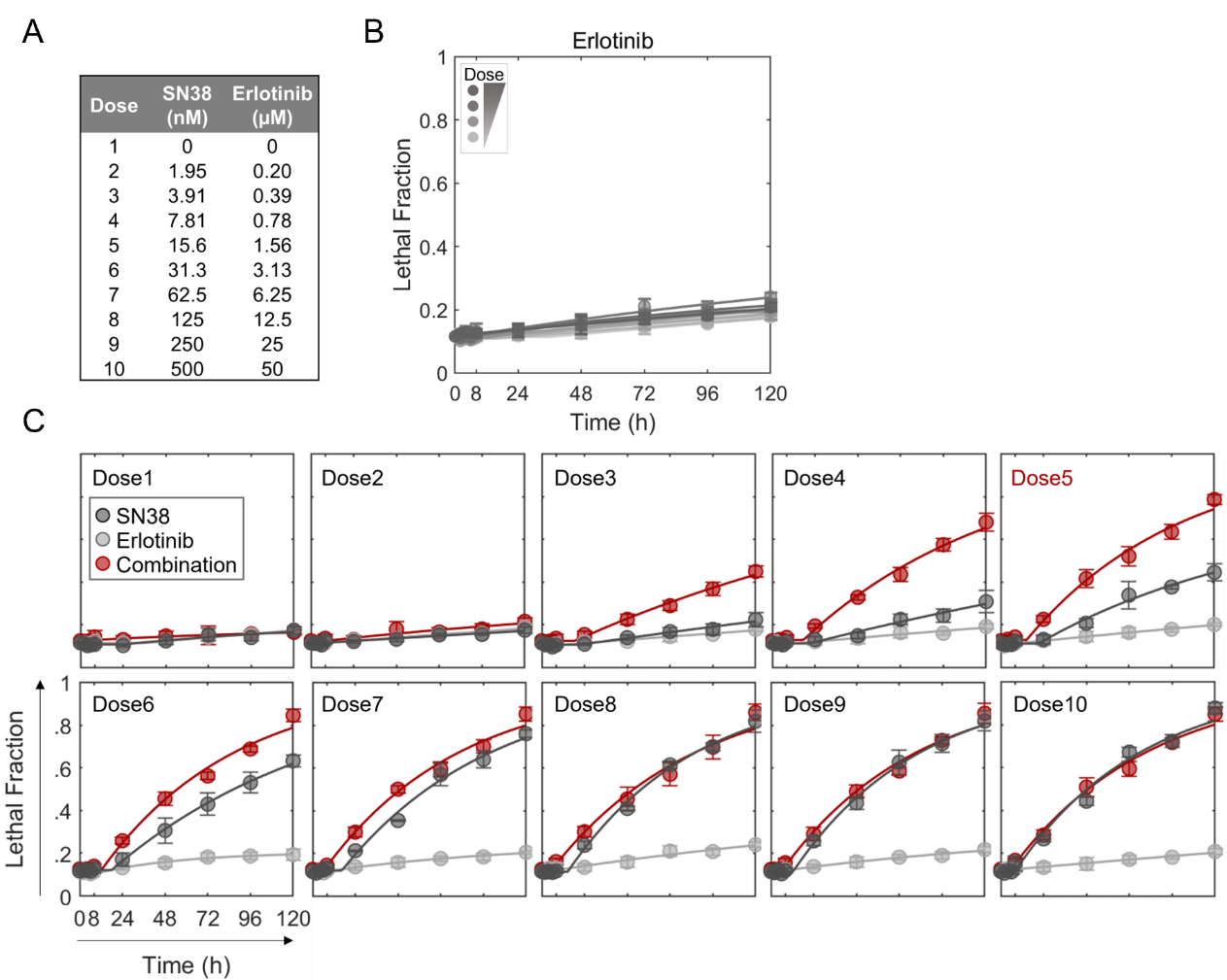

**Figure S2. (A)** The relative doses of SN38 and erlotinib used in the FLICK assay presented in Figure 2B-D, 4A, and 7E. **(B)** Analysis of the lethal fraction of erlotinib alone for all doses tested. Treatment with erlotinib alone did not substantially induce cell death in SNU5 cells. **(C)** Comparison of lethal fraction curves of the SN38/erlotinib combination and single-agent treatments at each relative dose. Curves at relative dose 5 highlighted are presented in Figure 2E.

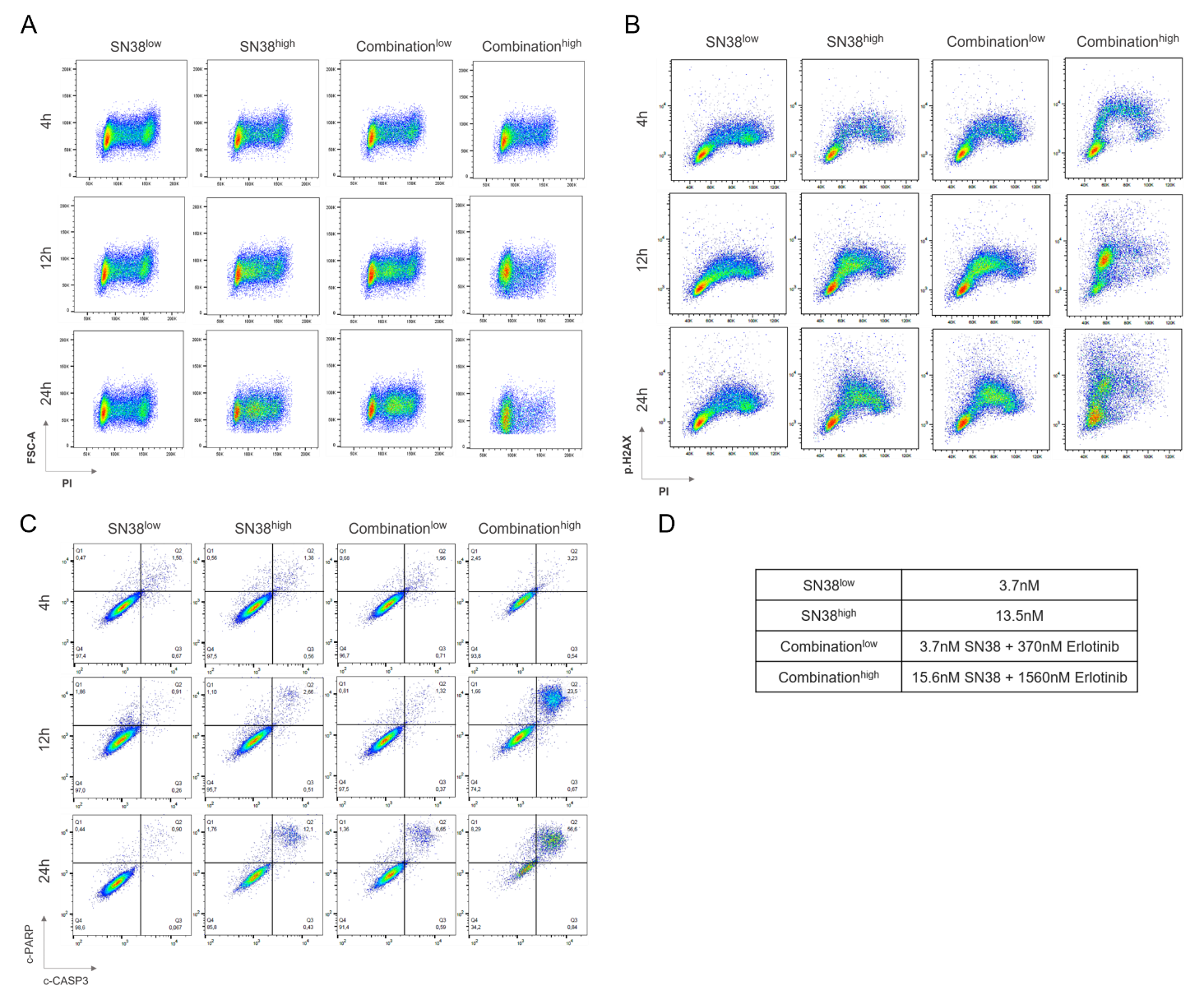

**Figure S3.** Representative flow cytometry plots for **(A)** cell cycle analysis, **(B)** DNA damage profiling coupled with cell cycle analysis, and **(C)** apoptotic cell death analysis in SNU5 cells. c-PARP: cleaved-PARP, c-CASP3: cleaved-caspase3, p.H2AX: phospho-H2AX, PI: propidium iodide. **(D)** Concentrations of erlotinib and SN38 used in (A-C).

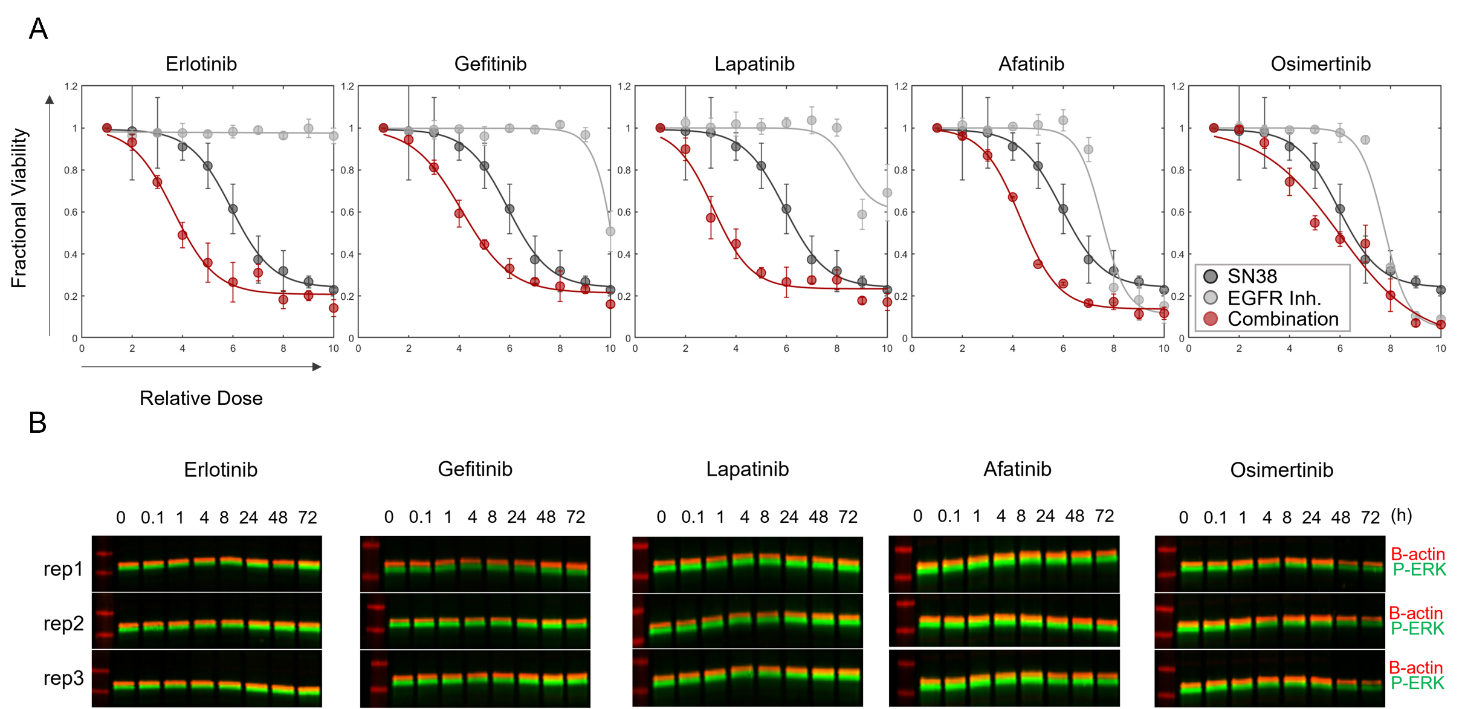

**Figure S4. (A)** Dose-fractional viability curves for the combination of SN38 with EGFR inhibitors erlotinib, gefitinib, lapatinib, afatinib, and osimertinib. **(B)** The immunoblots (as 3 biological replicates) to assess the effect of EGFR inhibitors on ERK phosphorylation.

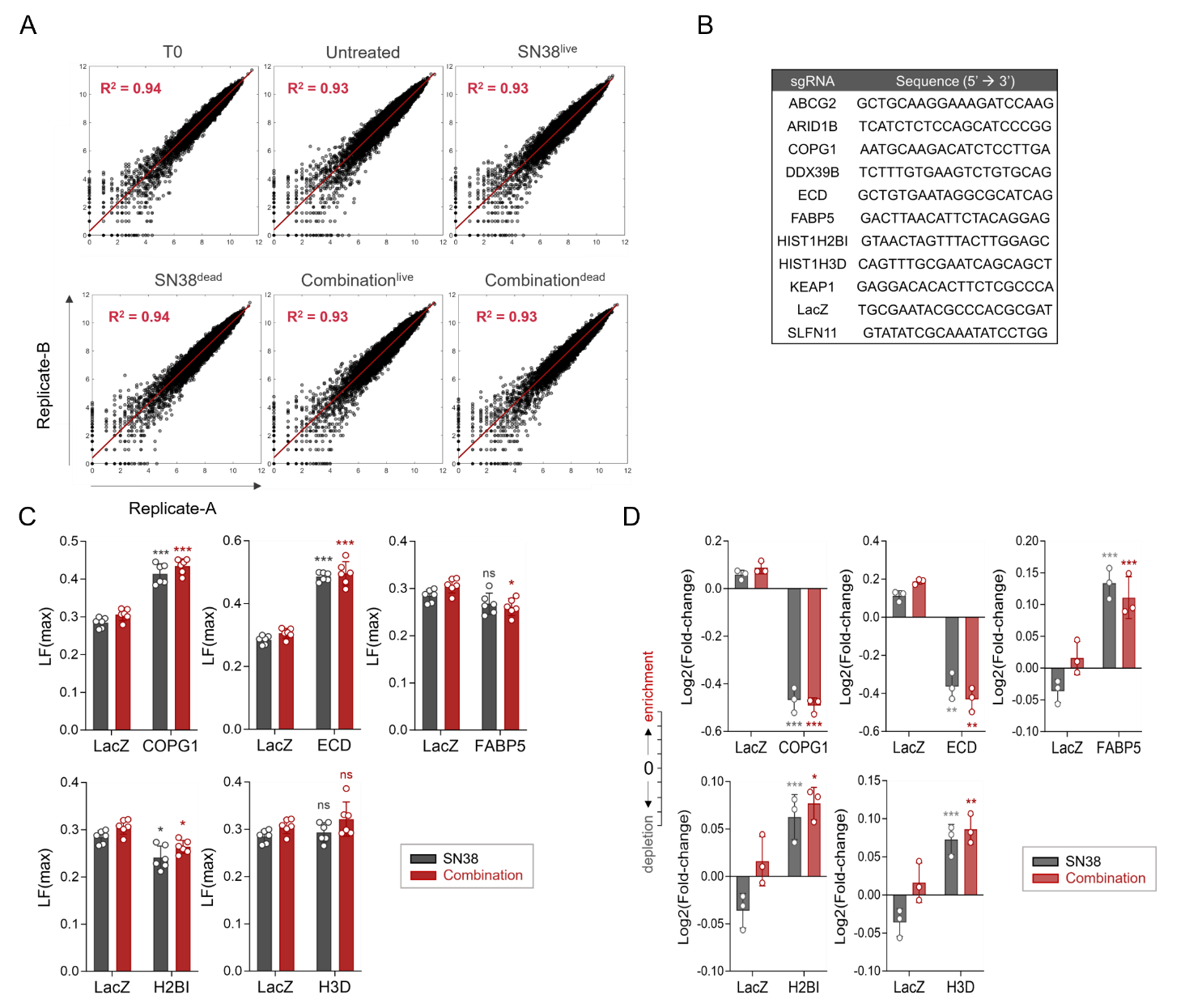

**Figure S5. (A)** Log_2_ count level correlation between the biological replicates of each experiment group. x-axis: replicate A at log_2_ count level. y-axis: replicate B at log_2_ count level. **(B)** sgRNA sequences used to validate the CRISPR screen hits. **(C)** LF (max), and **(D)** Log_2_ fold change plots for COPG1, ECD, FABP5, HIST1H2BI (H2BI), or HIST1H3D (H3D) knockout SNU5 cells or untargeted SNU5 cells (LacZ) under SN38 or SN38/erlotinib combination treatment.

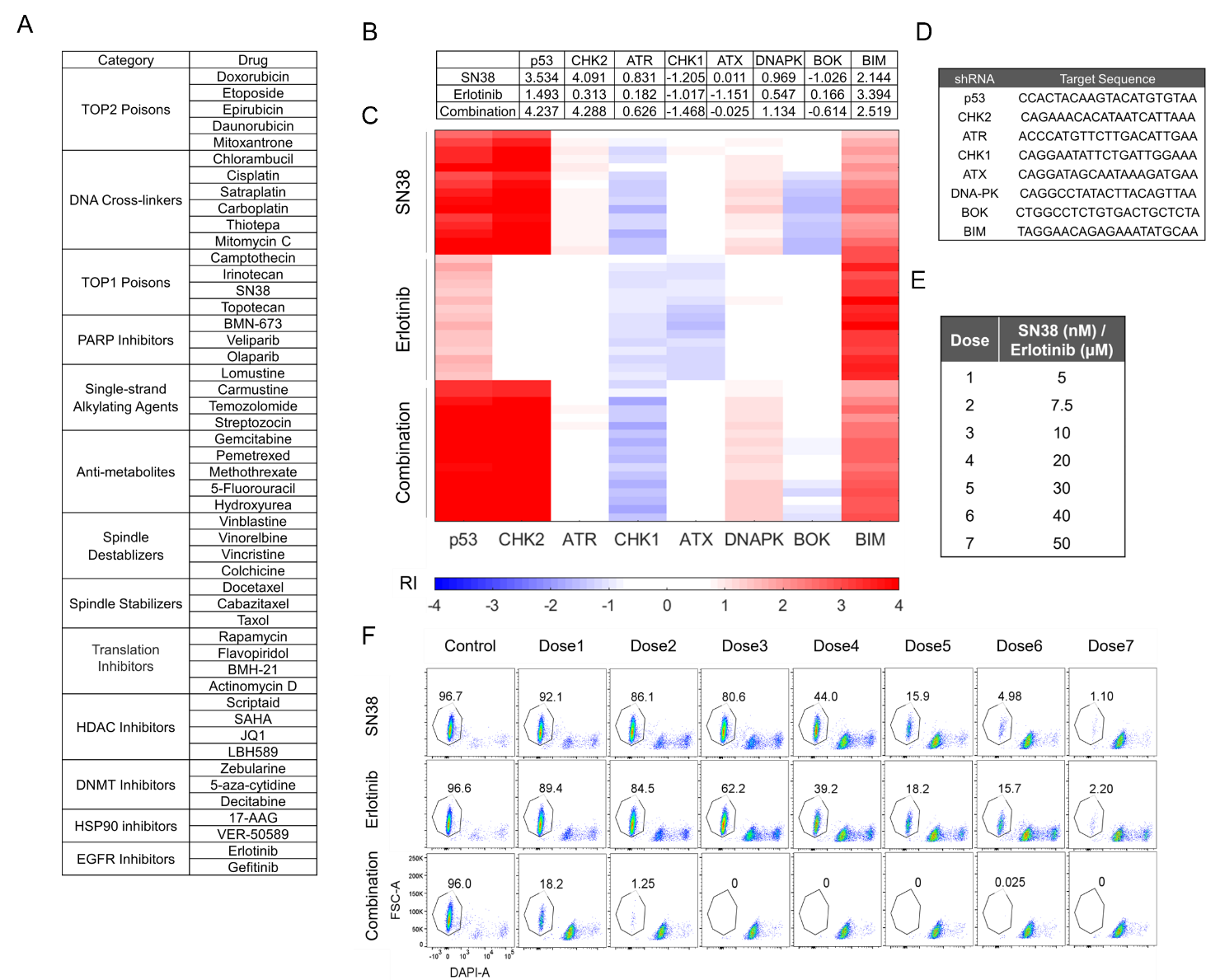

**Figure S6. (A)** The reference list of drug groups and drugs used to establish shRNA-based signature assay by Pritchard et al. **(B)** Average resistance index (RI) values for each cell population expressing the specified shRNA treated with SN38, erlotinib or SN38/erlotinib combination. **(C)** The heatmap of the signatures for all replicates of each treatment condition generated by assembling the RI values. **(D)** the shRNA sequences used in the signature assay. **(E)** The relative doses of SN38 and erlotinib applied for the dose-response analyses in Eμ-Myc Cdkn2a^Arf−/−^ cells, presented in Figure 6D. **(F)**Representative flow cytometry plots of live cell fractions for each tested dose of SN38, erlotinib, or SN38/erlotinib combination in the signature assay.

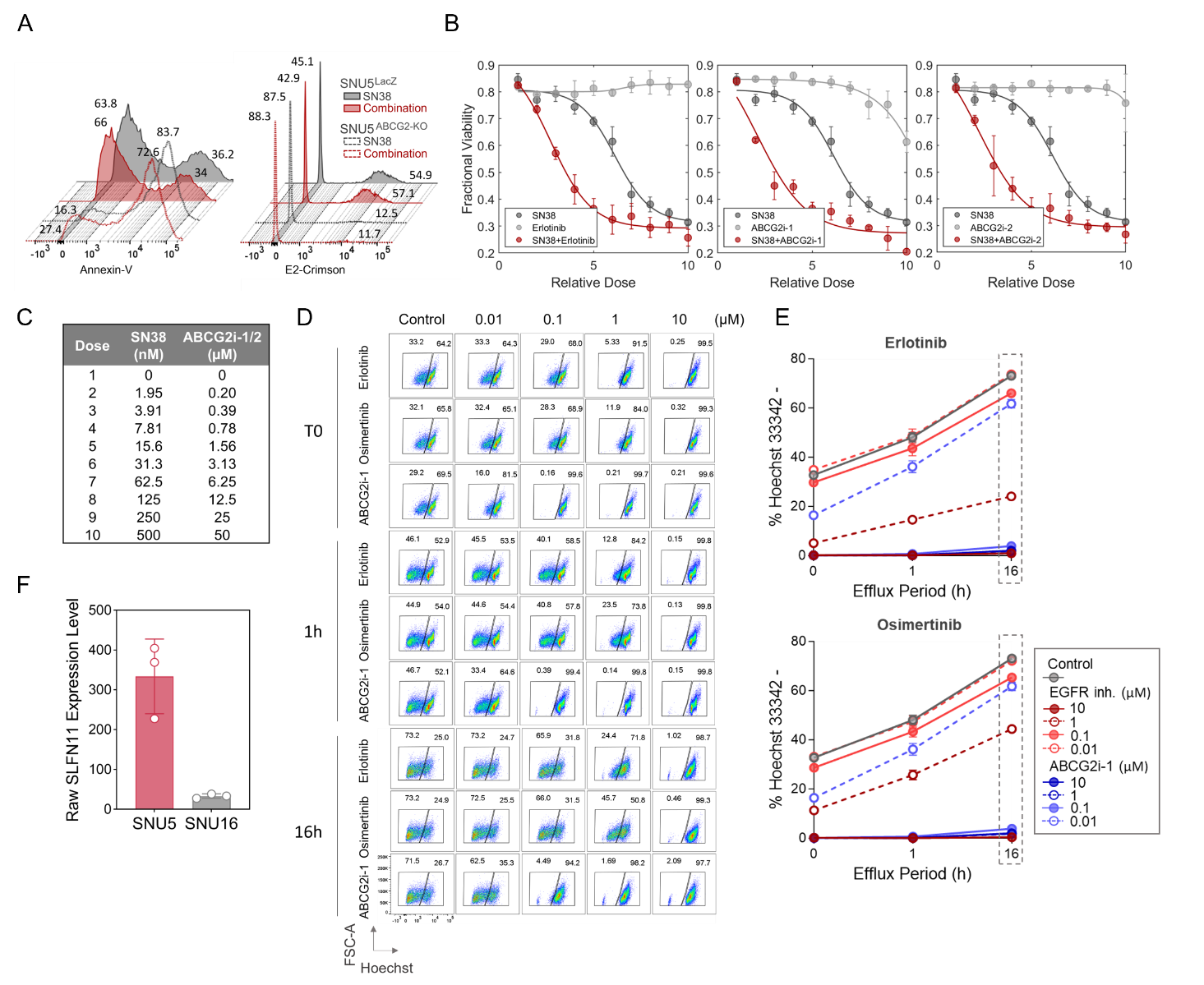

**Figure S7. (A)** Representative histogram plots of annexin-V (left) and e2-crimson (right) positivity in SNU5^LacZ^ and SNU5^ABCG2-KO^ cells treated with SN38 or SN38-erlotinib combination. **(B)** Dose-fractional viability curves for the dual combination of SN38 with erlotinib, ABCG2i-1, or ABCG2i-2 in SNU5 cells. **(C)** The concentrations of SN38 and ABCG2 inhibitors for the curves in B and Figure 7F. **(D)** Assessment of the impact of ABCG2 and EGFR inhibitors on the efflux of Hoechst over time in SNU5 cells via flow cytometry. **(E)** Percent inhibition of Hoechst efflux by ABCG2 and EGFR inhibitors over time in SNU5 cells calculated from the flow cytometry plots in D. **(F)** The gene expression level of SLFN11 in SNU5 and SNU16 cells. Expression data was exported from [merav.wi.mit.edu/](http://merav.wi.mit.edu/).
